# Skin metatranscriptomics reveals landscape of variation in microbial activity and gene expression across the human body

**DOI:** 10.1101/2024.12.02.626500

**Authors:** Minghao Chia, Amanda Ng Hui Qi, Aarthi Ravikrishnan, Ahmad Nazri Mohamed Naim, Stephen Wearne, John Common, Niranjan Nagarajan

## Abstract

The skin microbiome plays an important role in immune homeostasis and skin health, and yet our understanding of *in vivo* microbial gene activity is hindered by the lack of a robust, non-invasive protocol for metatranscriptomics across skin sites. Circumventing the challenges of low microbial biomass, host contamination, and RNA stability, we developed a clinically tractable skin metatranscriptomics workflow that provides high technical reproducibility of profiles (Pearson r>0.95), uniform coverage across gene bodies, and strong enrichment of microbial mRNAs (2.5-40×). Applying this protocol to a cohort of healthy adults (n=27) across five different skin sites (n=102, paired metatranscriptomes and metagenomes), identified a striking divergence between transcriptomic and genomic abundances, with *Staphylococcus* species and the skin fungi *Malassezia* having an outsized contribution to the metatranscriptomic landscape at most sites despite their modest representation in metagenomes. Species-level analysis showed skin site-specific enrichment of gene expression (e.g. increased levels of secreted fungal phospholipase C on cheeks relative to scalp), and revealed how key pathways were transcriptionally active *in vivo* (e.g. propionate and 4-aminobutyrate metabolism, potentially impacting skin barrier function). Gene-level analysis identified diverse antimicrobial genes transcribed by skin commensals *in situ*, including several uncharacterized bacteriocins, some of which are expressed at levels comparable to known antimicrobial genes. Correlation of microbial gene expression with organismal abundances uncovered >20 genes that putatively mediate interactions between microbes (e.g. a secreted *Malassezia restricta* protein with strongly negative *in vivo* association with *Cutibacterium acnes*; Spearman ρ>0.7). This work showcases the potential for leveraging skin metatranscriptomics to identify microbes whose activities play an outsized role in the community, and for uncovering pivotal microbial pathways and biomarkers linked to skin health and disease.

## Introduction

The human skin is home to diverse communities of microorganisms (bacteria, fungi, and viruses) that can interact with each other and the host to impact the skin microenvironment, immune homeostasis and skin health^1^. In recent years, our understanding of the role of the skin microbiome in various diseases has greatly benefited from the increasing accessibility of whole metagenome sequencing to identify key organisms and associated genetic potential differences^2–4^. This has been complemented by a growing number of *ex vivo*^5,6^ and *in vitro*^7,8^ studies that provide insights into mechanistic pathways through which the crosstalk between the host and the microbiome may be mediated. However, our understanding of whether these pathways are indeed utilized *in vivo* remains limited, as metagenomics only estimates functional potential of microbes^9^ (DNA content). Metagenomic DNA signals are a composite from living and dead cells, and even among living microbes, their genes can be variably expressed or transcriptionally silent in response to nutrient availability and other environmental cues^10–12^. Consequently, much remains unknown about which skin microbial species and pathways are transcriptionally active *in vivo*, their variability across skin sites and subjects, and their relationship to the skin metagenome.

Metatranscriptomics, which assays the pool of messenger RNAs (mRNAs) in a microbial community, has been used to study the transcriptional activity of microbiota in environments as diverse as the gut^13^ and ocean water^14^, but has not yet been broadly applied to the skin microbiome. Skin metatranscriptomics is hampered by the lack of a robust, non-invasive protocol that can accommodate a range of skin sites which have low microbial biomass, but substantial host and environmental contamination. This is because human skin is relatively sparsely colonized by microbes, with an estimated average density of 10^3^-10^4^ prokaryotes/cm^2^, several orders of magnitude lower than that of the gut^15^. Till date, there has been only one other published study which has leveraged RNA sequencing to reveal how microbes contribute to acne, albeit restricted to analyzing data from a single microbe (*Cutibacterium acnes*), and specifically applied for nose skin follicles using pore-stripping^16^. While biopsies are not limited to follicles and can capture more microbial biomass^17^, their invasive nature makes them impractical for adoption in large-scale clinical studies. Consequently, data from transcriptionally active microbes and microbial pathways across different skin sites is scarce and highlights the need for a generalized protocol to characterize skin metatranscriptomes^18^.

To address these challenges, we developed a robust workflow for non-invasive sampling and skin metatranscriptomics across body sites that demonstrates high technical reproducibility (Pearson r>0.95), uniform coverage across bacterial and fungal gene bodies, and strong enrichment of microbial mRNAs (2.5-40×). We further developed a data analysis workflow with rigorous control of “kitome” contaminants and taxonomic misclassification artifacts. The workflow is customized for profiling the skin metatranscriptome with the use of an updated skin microbial gene catalog based on metagenome-assembled genomes (MAGs) for diverse populations^19^. Leveraging this capability, we present the first multi-site metatranscriptomic survey of healthy human skin (n=27 subjects) from physiologically diverse skin sites (n=5; scalp, cheek, volar forearm, antecubital fossae and toe web). Analysis of paired metagenomes and metatranscriptomes (n=102 paired libraries; n=260 libraries in total) revealed a striking divergence between transcriptomic and genomic abundances, with *Staphylococcus* species and the skin fungi *Malassezia* having an outsized contribution to the metatranscriptomic landscape at most sites despite their limited representation in metagenomes. Species-level analysis showed skin site-specific enrichment of gene expression (e.g. increased levels of secreted fungal phospholipase C on cheeks relative to scalp), as well as revealed transcriptional activity *in vivo* of key pathways (e.g. propionate and 4-aminobutyrate metabolism, potentially impacting skin barrier function). Gene-level analysis identified diverse antimicrobial genes transcribed by skin commensals *in situ*, including several uncharacterized bacteriocins, some of which are expressed at levels comparable to known microbially produced antimicrobial genes. Correlation of microbial gene expression with organismal abundances uncovered >20 genes that putatively mediate interactions between microbes (e.g. a secreted *Malassezia restricta* protein with strongly negative *in vivo* association with *Cutibacterium acnes*; Spearman ρ>0.7). Overall, this work highlights the importance of metatranscriptomics for a holistic view of active species, expressed microbial functions, key pathways and microbe-microbe interactions occurring *in situ* on skin.

## Results

### Development of a robust skin metatranscriptomics workflow

We optimized an experimental workflow that is robust for metatranscriptomics across different skin sites by systematically and extensively testing different sampling tools (skin tapes versus swabs), lysis conditions and RNA purification techniques (**Supplementary Data 1**; **Methods**). Our final optimized protocol utilized skin swabs, sample collection in DNA/RNA shield, bead beating, rRNA depletion using a custom oligonucleotide mix and a direct-to-column TRIzol purification step (**Methods**). The robustness of our protocol was assessed with a pilot cohort consisting of both a cross-sectional group (n=24 samples) and a longitudinal group sampled across three consecutive days (n=45 samples), representing five distinct skin microenvironments: scalp (Sc), cheek (Ch), volar forearm (Vf), antecubital fossae (Ac) and toe web (Tw). A notably high proportion of metatranscriptomic libraries were successfully sequenced in this cohort (66/69=95%), enabling generation of a target million microbial reads per sample in most samples (>84%, median=2.2 million microbial reads, 0.66 Gbp; **Supplementary Data 1**; **Methods**). Depletion of rRNAs resulted in substantial enrichment (2.5-40×) of reads corresponding to non-rRNAs (median>79.5% of reads; **Figure 1A**). Analysis of the resulting reads showed that the corresponding skin metatranscriptomes were highly reproducible across different skin sites, with very strong consistency of species profiles (Sorensen similarity≥0.98; **Figure 1B**) as well as microbial gene expression (Pearson’s r≥0.99; **Figure 1C**) between technical replicates. Analysis of samples from the longitudinal group highlighted that metatranscriptomes can exhibit substantial temporal stability at the gene level (median Pearson’s r≥0.897 within individuals; **Figure 1C**), while being slightly more variable at the species level (median Sorensen similarity≥0.768; **Figure 1B**), with the temporal gene/species level variation being significantly lower than inter-individual variation at the same skin site (Wilcoxon p-value<10^-11^; **Figure 1B**-**C**).

**Figure 1:**
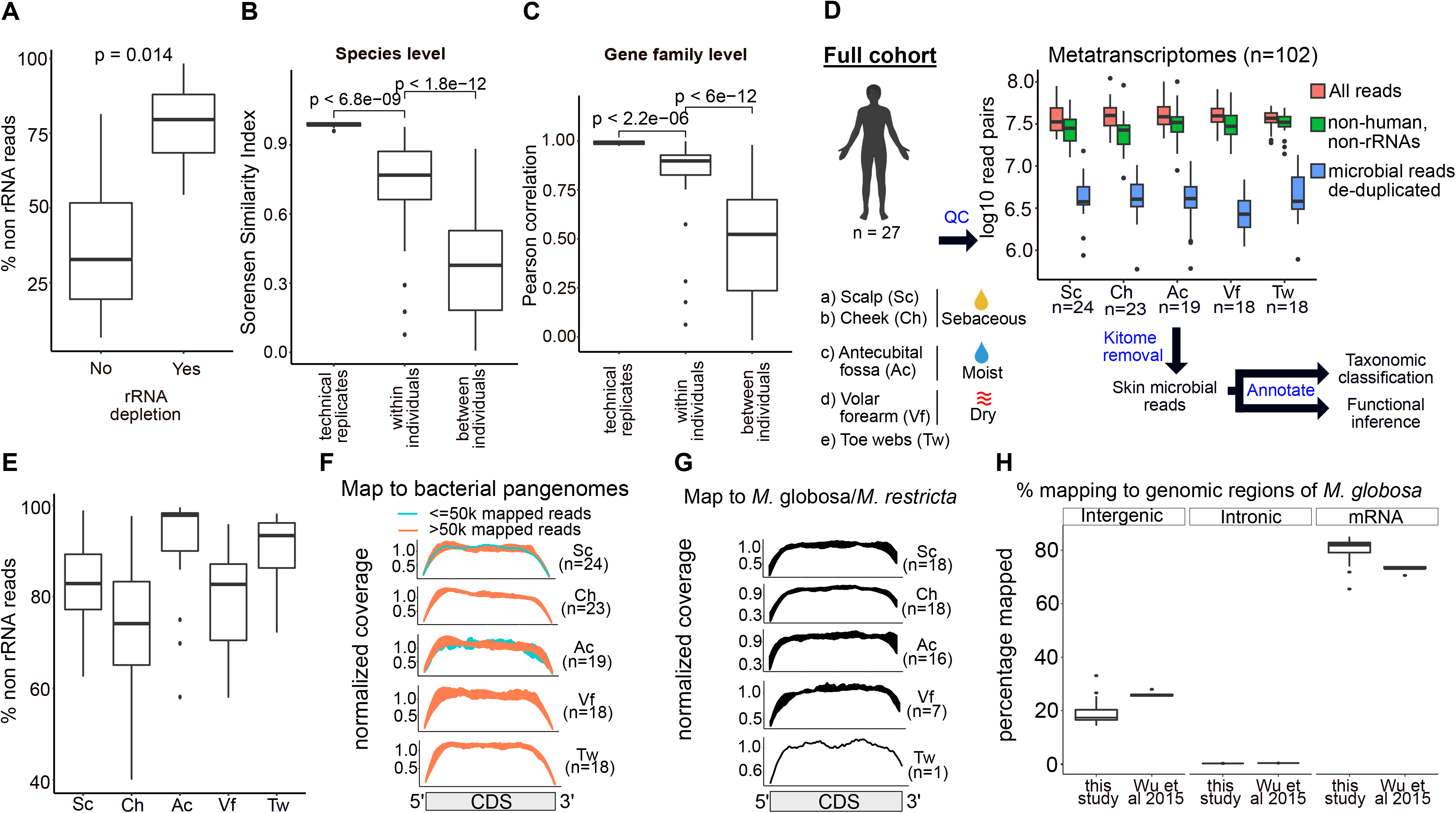
Robustness and reproducibility of skin metatranscriptomes across different skin sites. (A) Boxplot showing the fraction of non-ribosomal RNA (non rRNA) reads with and without experimental rRNA depletion during library preparation for the pilot cohort. (B) Boxplot of Sorensen Similarity Indices (1 – Bray Curtis dissimilarity) computed from species-level relative abundances for the pilot cohort. Pairwise similarities were computed between samples in three different categories. “Technical replicates” refer to different RNA-seq libraries prepared and sequenced from the same samples. “Within individuals” refer to data from re-sampling the same individuals across three consecutive days, while “between individuals” represents inter-individual variability in our cohort. (C) Same as B but showing Pearson correlation of gene expression signatures in the pilot cohort. (D) Schematic of the full cohort comprising 27 healthy adult volunteers and 5 different skin sites and the data analysis workflow. Boxplot showing the number of metatranscriptomic reads before computational removal of host and rRNA reads (red), number of non-human, non-rRNA reads after computational filtering (green) and number of microbial reads (comprising bacteria, viruses, fungi and archaea) after de-duplication (blue). (E) Boxplot showing the fraction of non-ribosomal RNA (non rRNA) reads with experimental rRNA depletion during library preparation for the full cohort. (F) Coverage plots showing the distribution of reads over bacterial gene bodies in samples belonging to the full cohort. Reads were mapped to bacterial pangenomes and samples with either >50,000 or ≤50,000 mapped bacterial reads are coloured differently. (G) Coverage plots showing the distribution of reads over fungal gene bodies for libraries (full cohort) with ≥500,000 fungal reads. (H) Percentage of reads mapped to different regions (intergenic, intronic and exons) of the *Malassezia globosa* genome. “This study” refers to samples from the full cohort. “Wu et al 2015” refers to an external RNA-seq dataset from cultured *Malassezia globosa* isolates^27^.

We next developed a computational workflow that could annotate skin metatranscriptomic reads with high sensitivity, while accounting for potential contamination signals and off-target matches. We firstly noted that using a skin-specific microbial gene catalog^19^ (IHSMGC) and a custom workflow, as opposed to a widely used general-purpose workflow (HUMAnN3^20^), resulted in a significantly higher median percentage of reads being functionally annotated (81% vs 60%, Wilcoxon p-value<3.1×10^-5^; **Supplementary** Figure 1A; **Methods**). We then used data from negative controls as well as prior reports^21^ to systematically identify potential DNA contaminant taxa^22^ (i.e. the kitome) and filtered corresponding reads from our analysis (**Supplementary** Figure 2A; **Methods**). Correlation analysis confirmed that skin microbes and potential kitome taxa (e.g. *Achromobacter, Bradyrhizobium, Mycolibacterium, Mycobacterium* and *Brevundimonas* species) formed distinct clusters of co-occurring taxa (**Supplementary** Figure 2B). To account for potential taxonomic classification errors, particularly in low complexity or misassembled regions of microbial genomes^23^, we compared a measure of unique matches in the genome based on minimizers with the total read count^23,24^ to identify false-positive taxa (**Methods**). Using data from spike-ins we found that an empirically determined threshold of unique minimizers per million microbial reads could discriminate false from true positive taxa at relative abundances as low as 0.1%, over a range of read counts (10^4^-10^6^ reads; **Supplementary** Figure 2C; **Methods**) and was therefore consistently applied as a filter.

We applied the combined experimental and computational skin metatranscriptomics workflow to the full cohort of 27 healthy individuals at 5 skin sites collected specifically for this study to provide comprehensive *in vivo* characterization of microbial gene expression on skin (n=135 paired samples for metagenomics and metatranscriptomics; **Figure 1D**, **Supplementary Data 1; Methods**). In the full cohort, the success rate for metatranscriptomic (102/135=75%) and metagenomic (130/135=96%) libraries was moderate-to-high, emphasizing the robustness of this protocol across a group of individuals. Typically, >1Gbp of de-duplicated non-rRNA sequencing data (median of 3.7 million read pairs) was generated per library similar to other metatranscriptomic studies^13,25^, with a relatively high median RNA quality across skin sites (DV200≥76; **Figure 1D**, **Supplementary** Figure 1B). In addition, paired metagenomes were sequenced to sufficient depth to obtain a median of 7 million microbial read pairs after filtering for human reads (**Supplementary** Figure 1C). Rarefaction analysis confirmed that libraries in the full cohort were typically sequenced at sufficient depths for representing active microbial functions/orthologous groups (>1 million read pairs; **Supplementary** Figure 3). Interestingly, the proportion of non-human reads was found to be significantly higher in metatranscriptomes versus metagenomes (98% vs 10%, Wilcoxon signed rank p-value<0.05; **Supplementary** Figure 1D), underscoring the feasibility of skin metatranscriptomic sequencing. Microbial rRNAs across all the sites were effectively depleted (2-25% of remaining RNA compared to 80-90% in a typical cell^26^) during the library preparation process (**Figure 1E**). In addition, the sequenced reads exhibited relatively even coverage across bacterial (**Figure 1F**) as well as fungal (**Figure 1G**) gene bodies in all five skin sites, and across a range of metatranscriptomic read depths. Similar to the pilot cohort, annotation rates for reads were found to be moderate-to-high across various body sites (median=69-80%, **Supplementary** Figure 1E). In addition, most of the reads mapping to the genome of the common skin fungi *Malassezia globosa* were in coding regions of mRNAs (>80%), with a minority mapping to intergenic regions and introns, indicating that our metatranscriptome libraries achieved DNA depletion levels similar to other high-quality *M. globosa* RNA-seq datasets derived from *in vitro* cultures^27^ (**Figure 1H**). Overall, these results emphasize that the generalized skin metatranscriptomics protocol presented here enables robust and reproducible profiling of microbial mRNAs across a wide spectrum of body sites and subjects.

### Skin metatranscriptomics identifies niche-specific active species and functions distinct from metagenomes

Different skin microenvironments (e.g. sebaceous, moist, or dry) greatly influence the composition of microbes that are present^4^, but much remains unknown about species or gene activities *in vivo*. While some studies have reported discordance between RNA and DNA abundances for specific species and gene families in gut and environmental microbiomes^14,25^, the scale and extent to which gene abundances and transcript levels correlate in the skin microbiome is not known. In this context, we observed a striking disparity between the most active species in skin metatranscriptomes versus the most highly abundant species in skin metagenomes (**Figure 2A**). For example, *Cutibacterium acnes* which is a dominant component of the metagenome at most skin sites (46-90% median relative abundance; except toe webs) has a relatively modest contribution to the metatranscriptome (2-31% median relative abundance). In contrast, while the skin fungi *Malassezia restricta* and *Malassezia globosa* were present at relatively low metagenomic abundances relative to bacteria (3-8% and 0.1-12% median relative abundance, respectively), they contribute substantially to metatranscriptomes in a niche dependent manner. *M. restricta* RNAs are heavily represented in cheek and scalp sites (23-30% median relative abundance), whereas *M. globosa* RNAs predominate on the scalp, antecubital fossae and volar forearms (21-81% median relative abundance). Toe webs were the exception to this observation, with a distinct microbial composition dominated by *Staphylococcus hominis* and *Staphylococcus epidermidis* in metagenomes and metatranscriptomes (**Figure 2A**). While normalized RNA counts of many commonly reported skin microbes varied positively with genomic DNA counts, several species had relatively large differences (≥4-fold) in RNA and DNA abundances, with disproportionately higher contributions to transcriptional activity (**Supplementary** Figure 4). *M. restricta* and *M. globosa*’s outsized contribution to the active biomass across different skin sites is likely driven by the larger cell volume of eukaryotes compared to prokaryotes^28^. Interestingly, *Staphylococcus* and *Corynebacterium* species also contributed proportionally more RNAs at the scalp, cheek, volar forearm and antecubital fossa after accounting for their metagenomic abundances (**Supplementary** Figure 4). The opposite was true for *C. acnes* and *Micrococcus luteus*, likely reflecting a relatively low proportion of cells present that are transcriptionally active. Overall, the stark differences between DNA and RNA abundances for various skin microbes underlines the need for skin metatranscriptomic measurements to characterize *in vivo* microbial activity.

**Figure 2:**
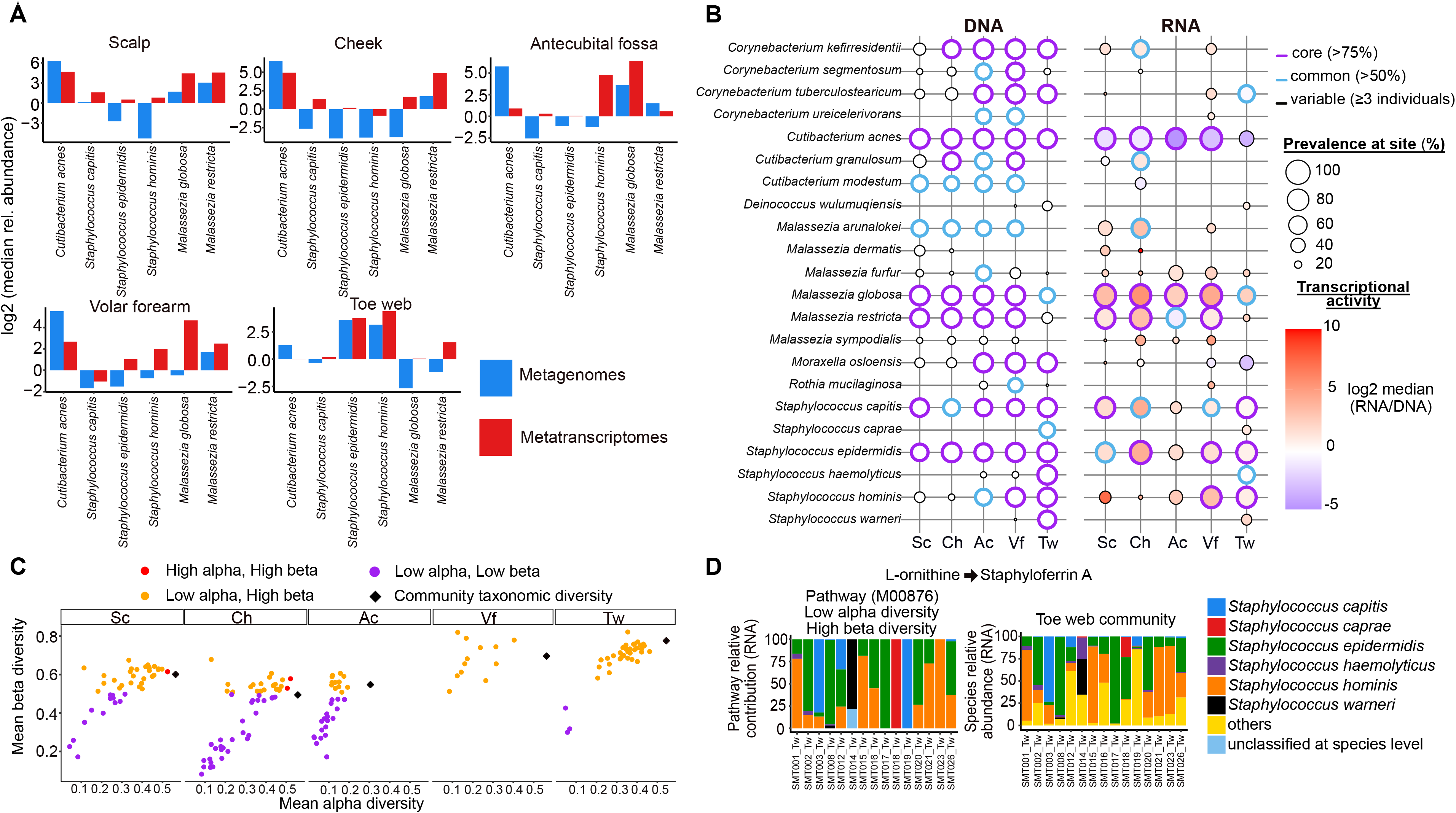
Niche specific signatures of core active species and transcribed functions. (A) Bar plots showing median relative abundances (total sum scaled) of reads assigned to various skin commensals across different skin sites presented in log_2_ scale. Abundances for metagenomes and metatranscriptomes are shown. (B) Bubble plot showing core, common and variable components of skin metagenomes (DNA) and metatranscriptomes (RNA) across different body sites. A species was called present in a metagenome if it had ≥0.1% relative abundance. A species was called present in a metatranscriptome if it had ≥0.1% relative abundance and was also detected in the metagenome. Core species in metagenomes and metatranscriptomes were defined as those present across >75% of samples at a given skin site. Common species for a skin site were defined as those present between 50% and 75% of samples. Variable species for a given skin site refer to other species that do not fall into the previous two categories, but which are present in ≥3 individuals. For RNA, bubbles are also shaded according to median transcriptional activity, defined as the ratio of normalized RNA counts of a species to that of their DNA. For visual clarity, only species which were core in at least one skin site are represented here. (C) Scatterplot of mean beta diversity (1 - Bray-Curtis dissimilarity) against mean alpha diversity (Simpson index) of core microbial pathways. Core microbial pathway expression was computed using HUMAnN3. Core microbial pathways were defined as those which were present (non-zero expression) at a skin site in >75% of individuals and with <25% unclassified reads at species-level. (D) Stacked bar plots for species level pathway contributions at RNA level, estimated with HUMAnN3 for Staphyloferrin A biosynthesis, and with Kraken2 for community level relative abundances at RNA level. For all sub-figures, Sc, Ch, Ac, Vf and Tw indicate the skin sites scalp, cheek, antecubital fossa, volar forearm and toe web respectively.

To profile the landscape of skin microbial activity across different niches, we categorized skin microbial species in terms of their prevalence as belonging to ‘core’ (present in >75% of samples), ‘common’ (present in 50-75% of samples) and ‘variable’ (present in <50% of samples) components of metagenomes and metatranscriptomes at each site. The median ratio of RNA/DNA levels per species was used to quantify transcriptional activity while accounting for variations in genomic abundances and sequencing depth (**Figure 2B**). Several species were found to be core in the metagenomes of all or most surveyed skin sites such as *Corynebacterium kefirresidentii*, *C. acnes*, *M. globosa*, *M. restricta*, *Staphyloccocus capitis* and *S. epidermidis* (**Figure 2B**). However fewer species were core components of the skin metatranscriptome across sites (e.g. *C. acnes*, *M. globosa* and *M. restricta*), with *Staphylococcus* species in particular exhibiting site-specific activity (e.g. *S. epidermidis* on cheek with median activity >17, *S. hominis* on volar forearm with median activity >9.5, and *S. capitis* on scalp with median activity >3.5), and variable prevalence across sites (e.g. in antecubital fossae; **Figure 2B**).

The same species also exhibited site-specific differences in overall gene expression activity that could reflect their preferences for certain skin microenvironments. Microbes that were prevalently found across multiple skin sites such as *C. acnes*, *S. capitis* and *S. epidermidis* showed higher overall transcriptional activity in sebaceous (scalp and cheek) versus non-sebaceous (antecubital fossae, volar forearm and toe web) sites, reflecting a general preference for environments richer in host lipids^29,30^ (**Supplementary** Figure 5). Intriguingly, although *M. globosa* and *M. restricta* are closely related lipophilic fungi, the latter showed larger variation in transcriptional activity between sebaceous and non-sebaceous sites, indicating that *M. restricta* may have greater sensitivity to host lipid availability (**Figure 2B**, **Supplementary** Figure 6). This is also in line with other reports demonstrating that *M. globosa* can be readily isolated from facial skin, the upper trunk and arms, while *M. restricta* recovery is mostly restricted to the scalp and face^31^. Within skin sites, individual heterogeneity is also evident in the species that are dominant in the metatranscriptome. For instance, while transcripts from *Malassezia* spp. make up most of the metatranscriptomes of the antecubital fossae (median relative abundance 98%), two individuals exhibited metatranscriptomes primarily dominated by *S. hominis* and *C. acnes* respectively (SMT003_Ac and SMT011_Ac; **Supplementary** Figure 7, 3rd and 8th column from the left). These observations highlight that the distribution of active skin microbes is shaped by skin site-specific differences but also individual-specific variations in the microenvironment.

To determine what functions are relatively important for microbial communities in distinct niches, we performed differential expression analysis of microbial genes clustered into orthologous groups (OGs), accounting for variations in DNA abundances (**Methods**). Compared to those on a sebum rich environment such as the cheek, bacteria colonizing the relatively dry volar forearms were found to upregulate gene clusters important for glucose catabolism, energy generation and protein synthesis, reflecting differences in resource availability (**Supplementary** Figure 8A**, Supplementary** Figure 9A). *Malassezia* fungi colonizing the cheeks upregulated gene clusters involved in mitotic growth, aromatic compound biosynthesis and protein modification relative to those in the volar forearms, consistent with increased lipid availability on the cheek for fungal growth and metabolism^28,32^ **(Supplementary** Figure 8A**, Supplementary** Figure 9A). Gene clusters involved in the citric acid cycle, amino acid metabolism, heme biosynthesis, RNA modification and kinase activities were upregulated in the toe webs relative to the volar forearms, indicative of microbes adapting their metabolism to an amino-acid rich environment there (**Supplementary** Figure 8B). Amino acid metabolic pathways are noteworthy because sweat is a rich source of free amino acids^33^ and the toe webs represent an environment high in sweat content relative to the volar forearm. The biosynthesis of heme, an important co-factor for many microbial processes involved in *Staphylococcal* colonization is also associated with increased availability of amino acids^34,35^. Consequently, most enzymes in the heme biosynthetic pathway from glutamate were upregulated by two-fold or more in toe webs compared to the volar forearm (**Supplementary** Figure 9B**-C**). Consistent with the more exposed nature of volar forearms and lower concentrations of protective secretions compared to cheeks^36^, microbes on volar forearms had upregulated expression of genes important for antioxidant protection relative to cheeks (**Supplementary** Figure 8A) and toe webs (**Supplementary** Figure 8B). These examples highlight the utility of metatranscriptomics for identifying key niche-specific functions in the skin microbiome.

We evaluated the importance of different species to core (present in >75% of individuals) biochemical pathways across different skin sites, as well as within-sample (alpha) and between-sample (beta) contributional diversities per pathway^9^ (**Methods**). At all sites besides the toe webs, *Malassezia* fungi were the predominant (>50% contribution) effectors of a substantial fraction (median 56%) of all core pathways compared to other bacterial genera (Wilcoxon p-value<0.05; **Supplementary** Figure 10A). Within bacteria genera, *Staphylococcus* was the major contributor to more core pathways than *Cutibacterium* at moist environments such as antecubital fossae and toe webs (Wilcoxon p-value<0.05; **Supplementary** Figure 10A). At any given site, some pathways might be redundantly expressed by multiple microbes while others might only be driven by one or a few species. Most core microbial pathways on skin were expressed by a few species within the same individual, but the same functions were expressed by diverse sets of species across individuals, indicating high functional plasticity in skin microbial communities (**Figure 2C**). For example, staphyloferrin A biosynthesis from L-ornithine is usually expressed by 1-2 *Staphylococcus* species in individual toe web communities but multiple possible *Staphylococcus* species can drive expression of this pathway across individuals (**Figure 2D**). Some metabolic functions were solely represented by fungi such as pathways for beta oxidation of very long chain fatty acids (VLCFA) being primarily expressed by *Malassezia* species on both sebaceous (scalp) and non-sebaceous sites (antecubital fossae) within and across individuals (**Supplementary** Figure 10B). Interestingly, other metabolic functions such as galactose degradation or arginine biosynthesis could be shared between skin bacteria and fungi, especially in nutrient-rich niches like the scalp or cheeks (**Supplementary** Figure 10C-D). While bacterial arginine has been described as a source of natural moisturizing factor (NMFs) for skin health^37^, our dataset indicates that fungi may be substantial contributors to the arginine pool. Overall, these results showcase how skin metatranscriptomics can be a complementary approach to metagenomics and demonstrates how different genera or species can act as the predominant effectors of niche-specific metabolic functions across skin sites.

### Species-level transcriptome analysis identifies signatures of microbial adaption to *in vivo* nutrient availability

Given the presence of niche-specific metatranscriptomic signatures, we next conducted species-level differential gene expression analyses to identify transcriptional variation in different environments that may highlight specific metabolic pathways essential for supporting *in vivo* colonization. We focused on RNA sequences belonging to the skin commensals *M. restricta*, *M. globosa, C. acnes* and *S. epidermidis*, because of their abundant transcript levels and importance for skin health^28,38^. In particular, *Malassezia* are highly represented in the metatranscriptomes of sebaceous sites (scalp and cheeks), while *S. epidermidis* can colonize both sebaceous and non-sebaceous sites (especially toe webs) and can be readily cultured *in vitro*. Differential expression analysis of *M. restricta* transcripts on scalp versus cheek showed that fungal genes and pathways associated with fructose and mannose metabolism, metabolism in diverse environments (e.g. *ACU7_1* and *acuE* of the glyoxylate cycle; *FBA1* and *KGD1* of the citric acid cycle; *TLK1* of the pentose phosphate pathway), and free fatty acid breakdown in peroxisomes (e.g. *POX2*, *POT1*, *PEX11B* and *ACOT8*) were upregulated on scalp relative to cheeks (**Figure 3A-B, Supplementary Data 2**). Conversely, genes and pathways associated with the breakdown of glycerides such as ether lipid and glycerophospholipid metabolism, as well as various secreted phospholipase C enzymes were relatively upregulated on cheeks (*PLC_1-7*; **Figure 3A-B**, **Supplementary Data 2**). This is consistent with distinct metabolite and lipid profiles on scalp and cheeks, with the former being relatively richer in nutritionally complex apocrine secretions and free fatty acids, while the latter being richer in glycerides^39,40^. Intriguingly, although *M. globosa* is another lipophilic skin fungi with comparable metatranscriptomic abundances to *M. restricta* on sebaceous sites (**Figure 2A**), there were few differences in gene expression between scalp and cheek sites for *M. globosa* (<30; **Figure 3C**). Another core skin commensal, *C. acnes*, had a similar number of differentially expressed features between the scalp and cheeks as *M. restricta*, but showed fewer significantly enriched pathways, with only those related to ribosome biogenesis, translation, and ATP generation being upregulated on the scalp (**Figure 3C**, **Supplementary Data 2**). These observations highlight how skin commensals may employ different gene expression strategies to thrive across body sites, including using specialized nutrient sources (*M. restricta*) and responding to energetic considerations (*C. acnes*).

**Figure 3:**
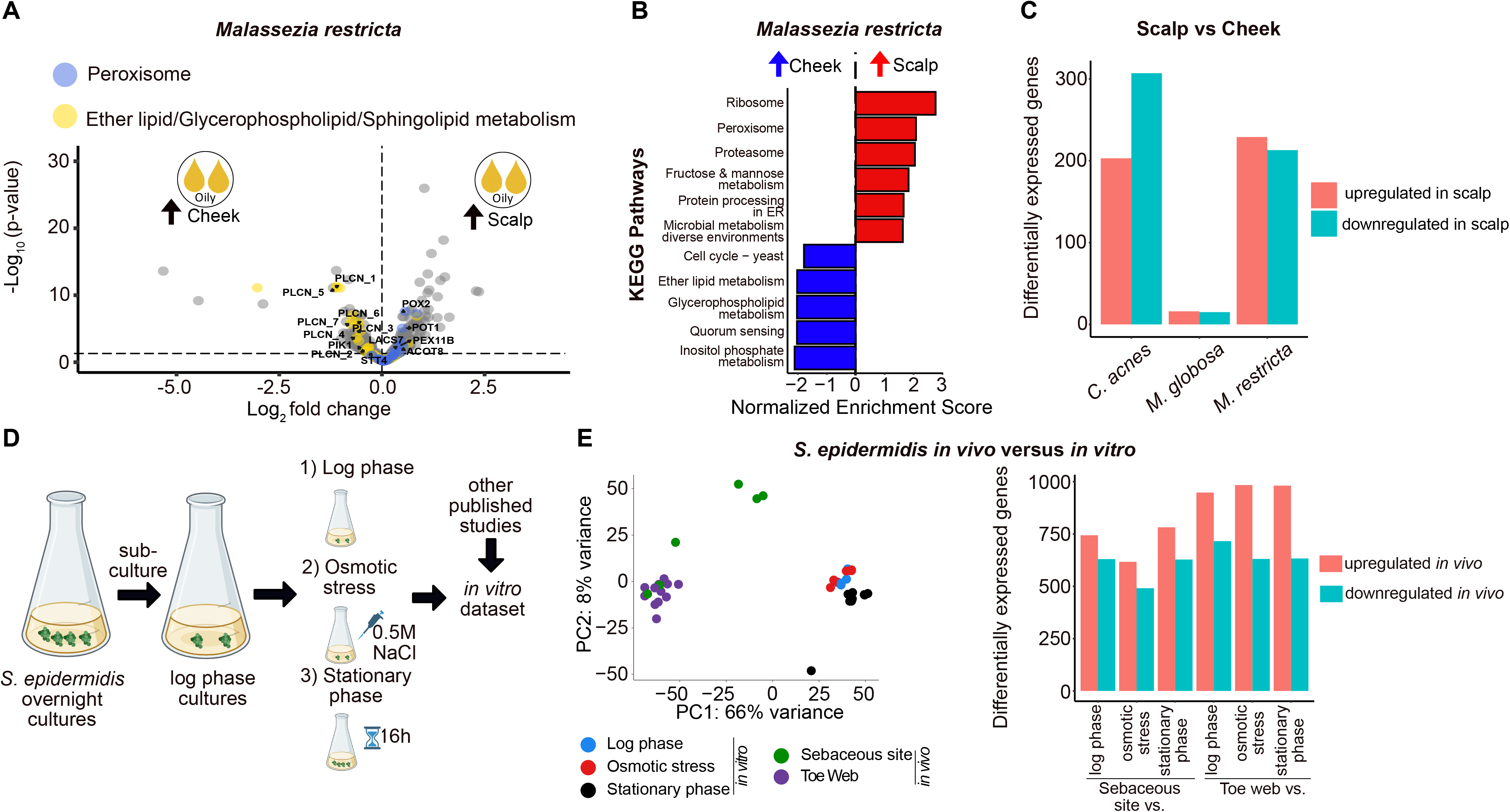
Species-level differential enrichment of metabolic pathways in various *in vivo* and *in vitro* growth conditions. (A) Volcano plot of differentially expressed genes for *Malassezia restricta* colonizing scalp versus cheek samples (≥200,000 *M. restricta* reads per library). Genes involved in peroxisomal activity or ether lipid/glycerophospholipid/sphingolipid metabolism are coloured blue or yellow respectively. The genes *PLCN_1-7* encode secreted phosopholipase C enzymes. The genes *POX2, POT1, PEX11B* and *ACOT8* encode proteins for peroxisomal function. (B) Bar plots of Normalized Enrichment Scores (NES) from Gene Set Enrichment Analysis for *Malassezia restricta* genes differentially expressed in scalp versus cheek samples. (C) Bar plots showing the number of differentially expressed genes between scalp and cheek sites for different species. (D) Schematic showing experimental setup for *in vitro* cultures of *Staphylococcus epidermidis* under different growth conditions and stress exposures. (E) Left: Principal component analysis (PCA) plot for *S. epidermidis* gene expression profiles from various *in vitro* and *in vivo* samples after batch correction using Limma. Right: Bar plots showing the number of differentially expressed genes for *S. epidermidis* between *in vivo* transcriptomes and transcriptomes for different *in vitro* conditions. Sebaceous sites refer to both scalp and cheek samples with ≥200,000 *S. epidermidis* reads per library (n=6). Toe webs refer to toe web samples with ≥200,000 *S. epidermidis* reads per library (n=12).

The transcriptomes of various skin microbes such as *S. epidermidis* have been well studied *in vitro*^41,42^ but not *in vivo*, raising important questions about how closely these culture models reflect their actual behaviour on human skin. Skin metatranscriptomics enabled the comparison of *S. epidermidis* gene expression *in vivo* versus three *in vitro* conditions comprising of cultures at log phase, stationary phase and subjected to osmotic stress, using the same RNA extraction and sequencing protocols (**Figure 3D**). Gene expression profiles between *in vivo* and *in vitro* conditions were clearly separated within the same experimental batch, or across different studies (generated in-house, Avican et al^42^ and Wang et al^41^) after batch correction (**Figure 3E left, Supplementary** Figure 11). There were many more differentially expressed genes between on-skin versus laboratory-grown conditions (n=1108-1664; **Figure 3E right**, **Supplementary Data 3, Supplementary Data 6; Methods**), relative to the modest changes between moist toe webs and sebaceous cheek/scalp sites *in vivo* (n=64 differentially expressed genes; **Supplementary Data 4**). Gene set enrichment analysis (GSEA) for *in vivo* vs *in vitro* comparisons revealed over 300 gene sets which were significantly enriched in either condition, with most differences observed between sebaceous sites and various *in vitro* conditions (**Supplementary** Figure 12, **Supplementary Data 5**; **Methods**). While *S. epidermidis* cultures *in vitro* upregulated genes in pathways primarily related to carbohydrate metabolism, reflecting the abundance of sugars in rich media, enriched pathways in sebaceous sites skewed towards sulphur metabolism, peptide and vitamin biosynthesis, reflective of the more complex nutritional landscape *in vivo* (**Supplementary** Figure 12). In contrast, there were far fewer significantly enriched gene sets/pathways in toe web sites relative to *in vitro* conditions despite having similar numbers of differentially expressed genes as sebaceous versus *in vitro* comparisons (**Figure 3E**, **Supplementary Data 3, Supplementary Data 6**). Nonetheless, sets of genes involved in metal ion homeostasis (GO:0030001) and copper ion binding (GO:0005507) were consistently upregulated in *S. epidermidis* colonizing toe webs relative to cultures at log phase or exposed to osmotic stress (**Supplementary** Figure 13A, **Supplementary Data 7**). These upregulated genes in the toe webs typically include pumps (e.g. efflux systems and P-type ATPases^43^) which couple ATP hydrolysis with the influx and efflux of different substrates and metal ions like copper and zinc (**Supplementary** Figure 13B). The upregulation of these pumps could be crucial for survival in toe webs, a “closed" environment rich in sweat containing trace metals excreted by the host^44,45^, where maintaining a fine balance of intracellular metal levels is essential for microbial cells^46^.

To contextualize overall differences in metabolic activity between *in vivo* and *in vitro* conditions via integration through metabolic networks, we performed flux balance analysis (FBA) based on genome-scale metabolic models constrained with our metatranscriptomic data^47^ (**Methods**). *S. epidermidis* showed distinct metabolic fluxes under all the conditions tested (83 metabolic reactions differing across all conditions, PERMANOVA Adonis R^2^=0.54, p-value<0.001), with a clear distinction observed for the *in vitro* conditions (log phase, osmotic stress and stationary phase) and even between *in vivo* conditions (sebaceous and toe web; **Supplementary** Figure 14A and **Supplementary Data 8)**. For example, fluxes associated with the production and export of propionate, a key short chain fatty acid impacting skin barrier function and immunity^48^, were higher *in vivo* for *S. epidermidis* compared to *in vitro* conditions (**Figure 4A**). Analysis of fluxes through glycolysis and pyruvate generation revealed that unlike under *in vitro* conditions, *S. epidermidis* predominantly generates the glycolytic intermediate glyceraldehyde-3-phosphate (G3P) via the pentose phosphate pathway (TALA, PFK_3) and the uptake of L-lactate (L_LACD2) to produce pyruvate for energy generation (**Figure 4B**). Pyruvate production *in vivo* is further supported by an active cataplerosis reaction (PPCKr), which generates intermediates contributing to pyruvate synthesis (**Figure 4B**). Under *in vivo* conditions, the NADH5 reaction which regenerates NAD, showed no flux (**Supplementary** Figure 14B). Instead, NAD regeneration might be driven by other compensatory cyclic reactions such as those involving 4-aminobutyrate consumption (ABUTR) and production (ABUTD) (**Supplementary** Figure 14C). While significant differences were identified in 58 reactions between the two *in vivo* conditions, overall *C. acnes* exhibited lesser variation in metabolic flux between the scalp and cheek (PERMANOVA Adonis R^2^=0.053, p-value<0.001; **Supplementary** Figure 15 and **Supplementary Data 9**). On both scalp and cheeks, *C. acnes* displayed an active Wood-Werkman cycle which facilitates propionic acid production^49^, with higher propionic acid production and export observed on the cheek compared to the scalp (GLM, adjusted p-value<0.001; **Figure 4C**). *In vivo* site-specific adaptations of *C. acnes* were also observed in the metabolism of specific amino acids. Flux analysis revealed that glutamate, a critical amino acid integral to linking key carbon and nitrogen pathways^50^, was produced via distinct metabolic routes in the scalp and cheek. On the scalp, glutamate production primarily occurred through histidine metabolism (**Figure 4D**), whereas on the cheek, it was synthesized predominantly through proline metabolism (**Figure 4D**), emphasizing niche-specific adaptation of metabolic pathways based on resource availability. Overall, these results highlight that species-specific analysis of metatranscriptomic data can be invaluable for identifying metabolic requirements for organismal survival on skin that would not otherwise be reflected in typical *in vitro* culture conditions.

**Figure 4:**
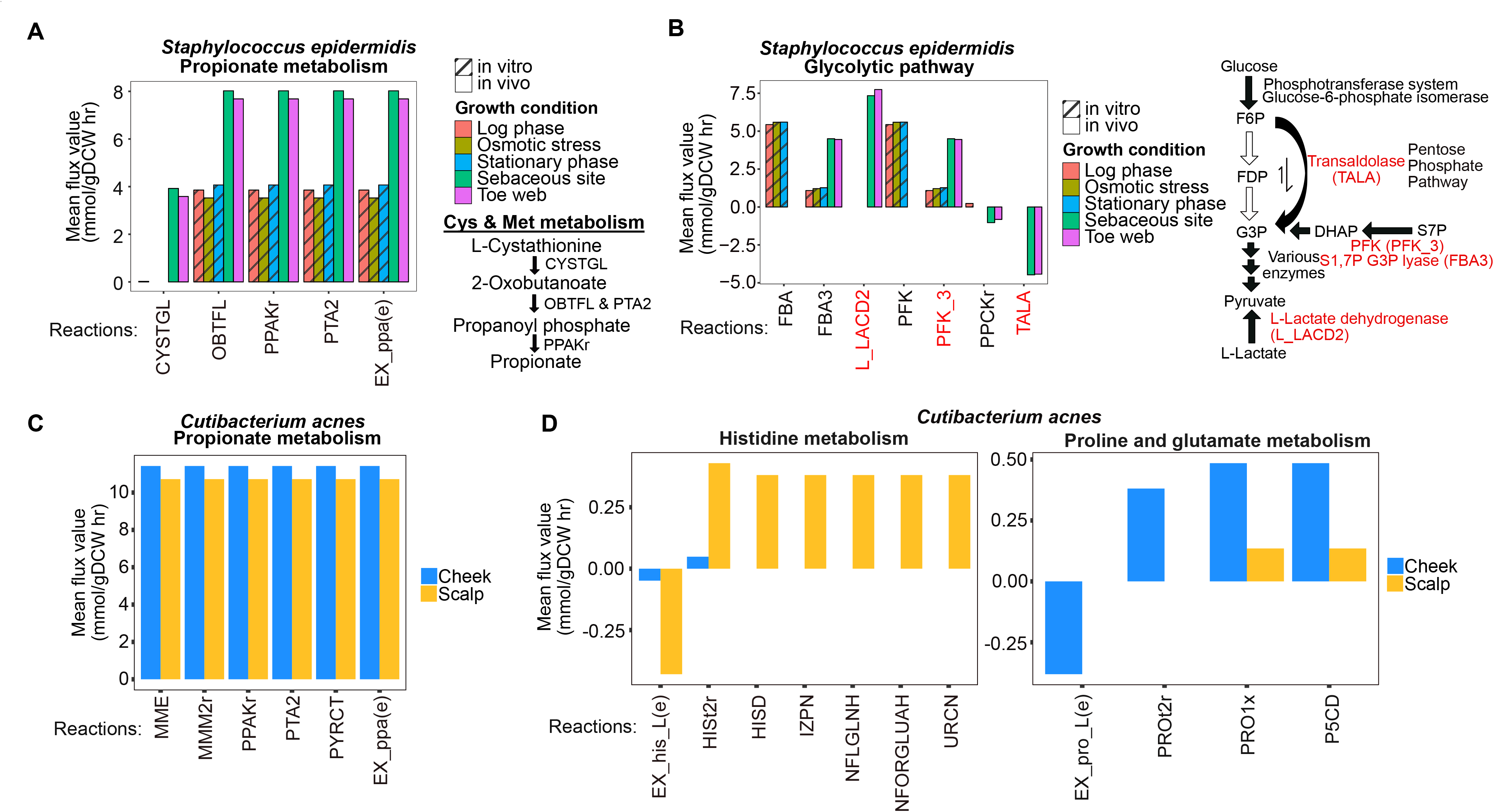
Transcriptome-aware, genome scale metabolic models reveal metabolic dependencies in various *in vivo* and *in vitro* growth conditions. (A) Bar plots showing mean flux values of various reactions associated with propionate production during cysteine and methionine metabolism in different *in vivo* and *in vitro* growth conditions for *Staphylococcus epidermidis*. Flux values were estimated using transcriptome-aware genome scale metabolic models for *S. epidermidis*. EX_ppae(e) represents the reaction exporting propionate out of bacterial cells. All reaction fluxes are significantly higher for *in vivo* relative to *in vitro* conditions (GLM adjusted p-value<0.001). (B) Same as A, except showing various reactions involved in pyruvate generation during glycolysis. All reaction fluxes are significantly different between *in vivo* and *in vitro* conditions. Reaction names highlighted in red are alternative means of generating pyruvate that have higher mean fluxes for *in vivo* versus *in vitro* conditions, consistent with *in vivo* metabolic dependency. (C) Bar plots showing mean flux values of various reactions associated with propionate production during cysteine and methionine metabolism in different *in vivo* growth conditions (Scalp or Cheek) for *Cutibacterium acnes*. Flux values were estimated using transcriptome-aware genome scale metabolic models for *C. acnes.* (D) Same as C, except showing various reactions involved in histidine and proline/glutamate metabolism. EX_his_L(e) and EX_pro_L(e) refer to transport reactions exporting histidine and proline respectively.

### Gene level analysis identifies key antimicrobial functions and interactions *in vivo*

As antimicrobial peptides and proteins can further shape niche adaptation and inter-species interactions, we searched for actively expressed genes that play a role in microbial competition *in vivo*. Antimicrobial peptides (AMPs) and proteins were sensitively detected by scoring “hit” database genes with profile hidden Markov Models (pHMMs; **Methods**). Skin microbes showed *in vivo* expression of a diversity of bacteriocins, phenol soluble modulins, enzymes that generate free radicals, and auto-inducing peptides (AIPs), representing classes of antimicrobial products that have been validated using *in vitro* or *ex vivo* models^7,8^ (**Figure 5A**). We also determined which species in our dataset expressed various classes of antimicrobial products (**Figure 5B**), motivated by the importance of identifying candidate species or bio-actives for antimicrobial therapies and further experimental follow-up. AIPs were expressed by different *Staphylococcus* species, consistent with the roles of these peptides in *Staphylococcal* quorum sensing and inter-staphylococcal competition^8^. Of particular interest are a diverse class of peptides called bacteriocins, which are secreted by bacteria or archaea to inhibit the growth or activities of other microbes in the community^51^. For example, some individuals were colonized by strains of *S. hominis* and *S. epidermidis* which expressed two peptides of the lacticin 481 family that were recently characterized to have broad antimicrobial activity against a range of gram-positive bacteria^52^ (**Figure 5B**). In contrast, relatively few individuals harboured *Staphylococcal* species with measurable expression of the well-characterized gallidermin/nisin family of bacteriocins which could be due to strain differences or its documented autotoxicity^53^.

**Figure 5:**
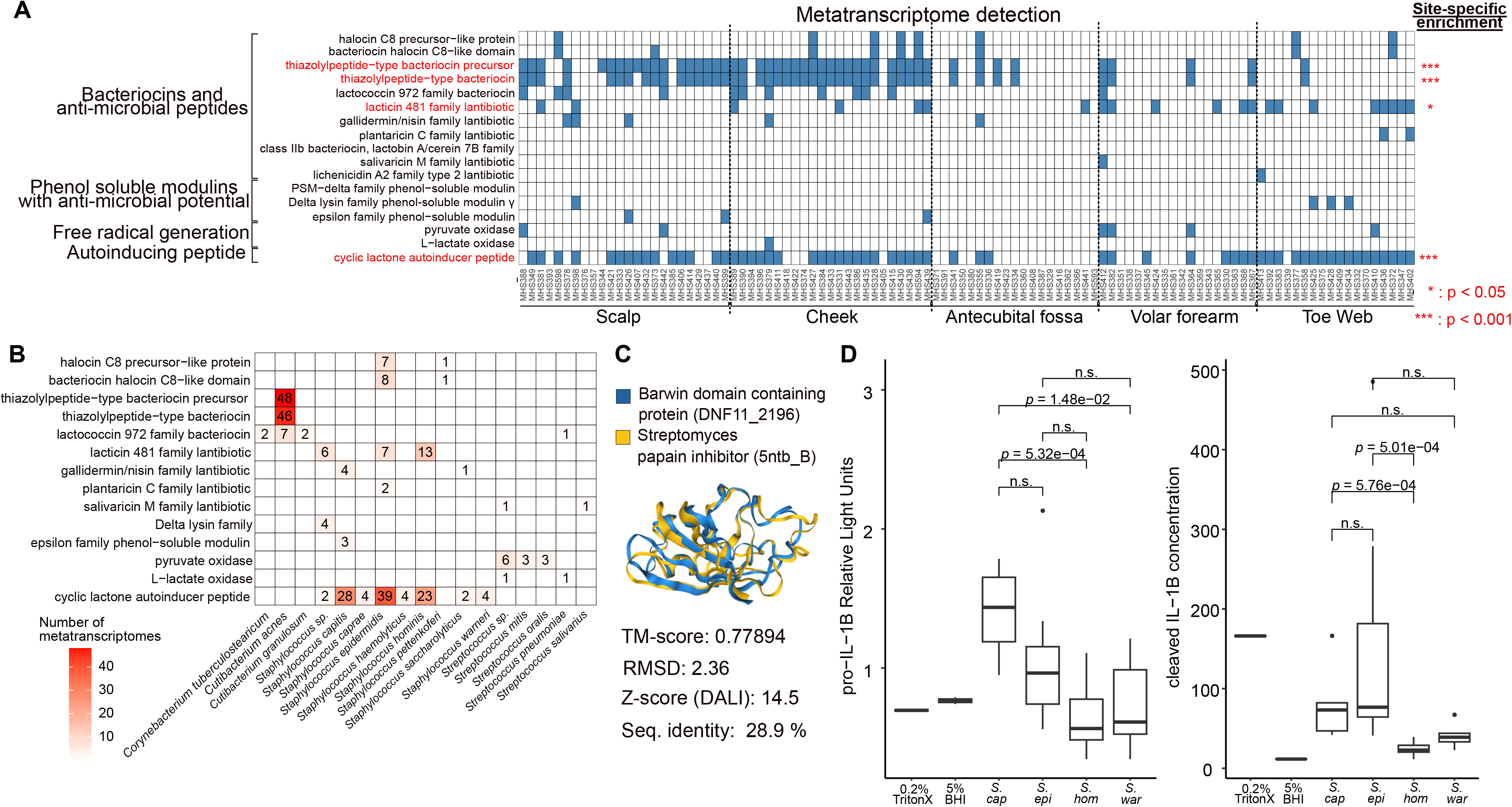
Inferring host-microbe and microbe-microbe interactions from metatranscriptomes. (A) Heatmap of various classes of antimicrobial genes with detected expression (≥5 reads) in individual metatranscriptomes grouped by skin site. Blue tiles denote detected expression in a given metatranscriptome. Red labels denote genes with site-specific enrichments based on Fisher’s exact test. Bacterial genes were grouped into categories of antimicrobial features based on information from profile Hidden Markov Models downloaded from NCBI. (B) Heatmap of various classes of antimicrobial genes with expression associated with species-level classified metatranscriptomic reads. The number of metatranscriptomes with detected expression (≥5 reads) is shown in the figure. (C) Overlaid protein structures of the *Malassezia restricta* protein DNF11_2196 and a *Streptomyces* papain inhibitor. Structures were obtained from the publicly available AlphaFold database. Structural similarity scores from Foldseek (TM-score and Root Mean Square Deviation) and Dali (Z-score) are shown. Primary amino acid sequence identity between the two proteins is also given. (D) Left: Boxplots of pro-IL-1B levels from human keratinocytes as measured by HiBiT and averaged across multiple strains of *Staphylococcus capitis* (n=4), *Staphylococcus epidermidis* (n=4), *Staphylococcus hominis* (n=8) and *Staphylococcus warneri* (n=4). Data are from two biological repeats, each comprising of three technical replicates. A Kruskal-Wallis test was conducted to assess whether levels differed significantly between groups (Kruskal-Wallis chi-squared=19.854, df=5, *p-*value<0.0014) and the adjusted p-values from post-hoc Dunn tests are shown. Right: Boxplots of cleaved IL-1B levels from human keratinocytes as measured by ELISA and averaged across multiple strains of *Staphylococcus capitis*, *Staphylococcus epidermidis*, *Staphylococcus hominis* and *Staphylococcus warneri*. Data are from two biological repeats, each comprising of three technical replicates. A Kruskal-Wallis test was conducted to assess whether levels differed significantly between groups (Kruskal-Wallis chi-squared=33.864, df=5, *p*-value<2.6e-06) and the adjusted p-values from post-hoc Dunn tests are shown.

There were notable examples of site-specific distribution of antimicrobial products. For example, transcripts belonging to cyclic lactone AIPs, thiazolylpeptides and their precursors were more frequently detected on sebaceous sites like the scalp and cheeks compared to other sites (Fisher’s exact test, adjusted p-value<0.001; **Figure 5A**). In some cases, site-specific expression was detected despite similar abundances of their host genomes in the community. For example, expression of the lacticin 481 family of lantibiotics by *S. hominis* and *S. epidermidis* was not evenly distributed across skin sites, with higher frequencies at the volar forearms and toe webs (Fisher’s exact test, adjusted p-value<0.05; **Figure 5A**). However, there were no statistically significant differences in metagenomic or total RNA abundances of these two species between lacticin 481 expressors versus non-expressors (Wilcoxon rank sum test, p-value>0.05; **Supplementary** Figure 16A-B). Such expression variability could be due to strain or environmentally driven differences. In contrast, thiopeptide expression on skin was associated with increased metagenomic and total RNA abundances of *C. acnes*, pointing to this species as an important source of thiopeptide AMPs (Wilcoxon rank sum test, p-value<0.05; **Supplementary** Figure 16C-D).

Several of the skin microbial bacteriocins detected in this study remain uncharacterized, representing an untapped source for development of new antimicrobials. Thiopeptides are important bio-actives that can shape skin microbial communities, e.g. cutimycin is secreted by *C. acnes* colonizing the hair follicles and can inhibit the growth of *Staphylococcus* species^2^. Two different thiopeptides (MET_03151623 and MET_02967399) were expressed by *C. acnes in vivo* (**Supplementary** Figure 17). While the thiopeptide MET_03151623 was identical to cutimycin, the other thiopeptide (MET_02967399) represents a putative bacteriocin with uncharacterized properties because it shared <50% amino acid identity with cutimycin and other members of the thiopeptide family (**Supplementary** Figure 17). Some *Cutibacterium and Corynebacterium* strains also expressed peptides of the lactococcin 972 family, but most of them had primary sequences which differed from those characterized in bacilli such as *Lactococcus* (<40% identity; **Figure 5B**, **Supplementary** Figure 18). Unexpectedly, several individuals had strains of *S. epidermidis* and *Staphylococcus pettenkoferi* colonizing the toe webs which expressed putative bacteriocins homologous to those of the halocin family (**Figure 5B**, **Supplementary** Figure 19). Previously, members of this family were only described in halophilic archaea and bacteria^54,55^. These examples highlight the unexplored landscape of antimicrobial mechanisms deployed by various skin microbial species and strains to thrive in their *in vivo* niches.

Finally, we leveraged paired metatranscriptomics and metagenomics to identify putative host-microbe and microbe-microbe interactions. Despite the relatively low fraction of human reads in our metatranscriptomic libraries (median <3%), a median of 0.8 million human reads per library was obtained, similar to the depth of sequencing commonly seen in single-cell transcriptomics studies^56,57^ (**Supplementary** Figure 20A). Assessing our swab-based metatranscriptomics libraries relative to previous biopsy-based studies^58,59^ highlighted that a substantial proportion of human reads were exonic (median 31% versus <15% in a previous biopsy-based study^59^) and the proportion of intergenic reads was low (median 18%) relative to transcriptomic data generated in-house from skin biopsies (median 32%; **Supplementary** Figure 20B). Gene Set Variation Analysis^60^ (GSVA) was used to estimate sample-specific immune pathway activities to find associations between host immunity markers and metagenomic abundances of different microbes. This analysis revealed three statistically significant relationships, all of which involved *Staphylococcus capitis* (FDR<0.25; **Supplementary Data 10**; **Methods**). The IL-6/JAK/STAT3 and Toll-like receptor signalling pathways which are involved in Th17 function^61^ and microbial detection by the immune system respectively, were positively correlated with *Staphylococcus capitis* abundances on cheeks (**Supplementary** Figure 21). This is consistent with observations showing that high *Staphylococcus capitis* abundances on skin are associated with an IL-17 dominated immune profile in skin disease^62^. Testing this hypothesis further, we noted that exposure to the supernatants of different strains of *S. capitis* led to consistently higher levels of both pro-IL-1B and cleaved IL-1B in human keratinocytes relative to the induction by other commensal *Staphylococcal* species, in line with our findings that *S. capitis* is associated with induction of specific immune responses on skin (post-hoc Dunn’s test, adjusted p-value<0.05; **Figure 5D**).

To identify putative microbe-microbe interactions, the abundances of transcripts encoding proteins entering the secretory pathway in one species were correlated with organismal abundances of another species within the same skin site (**Methods**). A total of 13 different combinations of common skin commensals were tested transcriptome-wide and >30 significant associations were identified (FDR<0.1; **Supplementary Data 10**). Notably, transcript abundances of a protein (DNF11_2196) predicted to enter the secretory pathway in *Malassezia restricta* were strongly negatively correlated with normalized *C. acnes* abundances on scalp (Spearman’s ρ<-0.7, adjusted p-value<0.05; **Supplementary** Figure 22A). DNF11_2196 is a poorly characterized gene (https://alphafold.ebi.ac.uk/entry/A0A3G2S5R5) with low primary sequence identity to the top hit in the PDB100 structure database (28.9% identity, 81.2% query sequence coverage; Foldseek web server, 3Di/AA mode). This makes it challenging to infer function based on sequence-based homology alone. However, the availability of accurate protein structure folding and searching algorithms enables functional inferences based on similarities in 3D structure^63,64^. There was greater similarity at the protein structural level indicated by both Foldseek (TM-Score=0.77894, RMSD=2.36) and Dali (Z-score=14.5) between DNF11_2196 and the structure of a *Streptomyces* papain inhibitor (5ntb-B), which has antimicrobial properties due to its inhibition of bacterial cysteine proteases^65^ (**Figure 5C**). Importantly, the negative correlation between DNF11_2196 and *C. acnes* abundances could not be explained by an inverse relationship between the organismal abundances of *M. restricta* and *C. acnes* (**Supplementary** Figure 22B-D). A similar analysis done in cheek samples showed that *Cutibacterium granulosum* abundances were positively correlated with expression levels of a *C. acnes* triacylglycerol lipase, independent of relationships between organismal abundances, indicating a potential symbiotic interaction and mechanism (Spearman’s ρ>0.7, adjusted p-value<0.05; **Supplementary** Figure 22E**-G**). Overall, these results showcase that skin metatranscriptomic and metagenomic data can be integrated to identify sets of expressed genes mediating host-microbe and microbe-microbe interactions *in vivo*, and thus help to prioritize candidate bioactive molecules or signalling pathways for further characterization and validation in model systems and clinical trials.

## Discussion

We report here the development of a robust experimental and computational workflow tailored for metatranscriptomics of diverse skin sites, circumventing the challenges of relatively low biomass by optimising the sampling, extraction, rRNA depletion, contamination removal, and functional classification steps (**Figure 1A-C**; **Supplementary** Figure 1-2). By applying this to a cohort of healthy individuals across multiple skin sites, we reveal the landscape of active species, microbial signatures of adaptation to their niches, as well as several key microbe-microbe and host-microbe interactions *in vivo*. We note that unlike human stool metatranscriptomes^13^ and skin metagenomes, skin metatranscriptomes of the stratum corneum are predominantly composed of microbial reads (>97% of all reads before de-duplication). Consequently, skin metatranscriptomes sequenced to relatively modest sequencing depths (5-10 million paired-end reads) can capture most active microbial functions at the community level based on rarefaction analysis (**Supplementary** Figure 3). The abundance of microbial reads means that skin metatranscriptomics can be feasibly deployed for population scale or longitudinal studies, especially as sequencing costs continue to decrease with newer platforms. Our protocol is compatible with a diversity of skin sites across individuals and non-invasive sampling using commercially available swabs makes deployment suitable for clinics to investigate disease cohorts. The moderate-to-high success rate of library construction from limited input, along with strong technical reproducibility and even coverage of mapped metatranscriptomic reads across microbial gene bodies, enables robust *in vivo* gene expression measurements and the assessment of changes in microbial activity in highly heterogeneous cohorts or over time. Given the very low biomass in the antecubital fossa and volar forearm relative to other skin sites, further experimental improvements could be explored to decrease the loss of nucleic acids during the extraction process, such as the use of depletable carrier RNAs and single-cell RNA-seq library kits that can tolerate lower inputs and hence increase robustness. Collapsing of read pairs with identical 5’ and 3’ ends is a conservative approach to de-duplication which removes a substantial proportion of microbial reads. More precise de-duplication methods based on unique molecular identifiers (UMIs) can be incorporated into this workflow to better distinguish PCR and biological duplicates^66^. While we optimized this workflow using swabs which sample the stratum corneum for their ease of use, our extraction and computational workflows are also suitable for profiling metatranscriptomes from deeper skin layers obtained from other sampling modalities such as biopsies, pore strips^16^ or follicular extracts^67^. Such studies will be necessary to enrich for signatures of immune cells which typically reside beneath the stratum corneum^68^, or for functional analysis of different *Cutibacterium acnes* strains which have been shown to colonize individual pores in a clonal fashion despite their co-existence across the skin surface^69^. Altogether, our workflow is a widely-applicable and clinically tractable approach to survey microbial activities on skin surfaces *in vivo*.

A key observation in our work is that skin metatranscriptomes yield distinct information about microbial activities compared to their matched metagenomes, similar to previously published studies in other contexts such as from human stool^13^ or ocean water^14^. Surprisingly, core microbial pathways on skin were expressed by relatively few species (low metatranscriptomic alpha diversity) in the community (**Figure 2C**), in contrast to stool metatranscriptomes which harboured numerous “housekeeping” pathways which were expressed by most species in the community^13^. This implies that unlike stool communities, a comparatively smaller fraction of microbes on skin surfaces are active, with the remainder being either quiescent or dead. Akin to our observations on skin, low metatranscriptomic alpha diversity of microbial pathways has also been previously described in nasal cavities and vaginal surfaces^9^, possibly reflecting nutrient scarcity in these environments which may support the robust growth of only a limited number of species. Of note, *Malassezia* and *Staphyloccocus* species had outsized contributions to skin metatranscriptomes, with disproportionally higher RNA abundances relative to their DNA. This can be attributed to a combination of factors such as cell size and bioactivity. As eukaryotic fungi, *Malassezia* cells have a volume and biomass that are at least two orders of magnitude greater than the average bacterial cell^28^. This means that metagenomes, which measure genome copy numbers, likely underestimate the contribution of *Malassezia* spp. to the functional potential of the skin microbiome. *Malassezia* species are important sources of secreted lipases, proteases and metabolites that can shape host and microbial activities on skin^7,70^. For example, *Malassezia* phospholipases can breakdown host lipids to generate polyunsaturated fatty acids (PUFAs) such as arachidonic acid which are potent inflammatory mediators^71^. The relatively high activity of various *Staphylococcus* species was unexpected, as they usually make up only a small proportion of bacterial communities on skin, except in sites such as toe webs. This could be reflective of the metabolic versatility of *Staphylococcus* spp. for survival in diverse skin microenvironments due to their ability to utilize a range of carbon sources, amino acids and lipids for their energetic and cellular demands^72–74^. Certain staphylococci such as *S. epidermidis* have also evolved a range of strategies to escape suppression by the host immune system such as intracellular localization^75^ or by expressing antigens that lead to commensal-specific T cell responses^76^. Our workflow will thus be a useful tool to study how *Staphylococcus* species contribute to skin health, especially given reported associations of these microbes with skin phenotypes such as malodor^77^, itch^6^ and eczema^10^.

Our work represents the first attempt to leverage metatranscriptomics to characterize the metabolic pathways required by different skin microbes to thrive in their *in vivo* niches. Besides identifying significantly enriched metabolic pathways computed from differential expression analysis, we further extended this approach by integrating transcriptomic data with flux balance analysis (FBA)^78^. The latter approach models organismal activity as a system of all predicted metabolic reactions instead of examining individual pathways in isolation, allowing us to identify metabolic requirements or dependencies necessary for maximizing biomass *in vivo* which cannot be readily determined from *in vitro* co-culture models^79^. Interestingly, while the lipophilic fungi *Malassezia restricta* was present in the metagenomes of both facial skin (cheeks) and scalp, metatranscriptomics revealed differential enrichment of fungal pathways that metabolize distinct classes of host lipids between the two sites. This observation highlights the importance of using metatranscriptomics to study organismal activities or phenotypes, even when there are no significant differences in organismal abundances. When applied to microbes growing under different *in vivo* versus *in vitro* conditions, transcriptome aware FBA revealed that the pentose phosphate pathway (PPP) and lactate metabolism were preferentially utilized by *S. epidermidis* to generate pyruvate as a respiratory substrate *in vivo* in contrast to *in vitro* conditions. This observation is consistent with previous reports showing that the PPP is crucial for energy production, biofilm formation and virulence in staphylococci^80^. Interestingly, our results also suggest that besides *Cutibacterium acnes*, commensal *S. epidermidis* may be another *in vivo* exporter of the short-chain fatty acid (SCFA) propionate, which has been linked to immunomodulation in keratinocyte and sebocyte cell lines^81^. Further studies are needed to determine if propionate mediates crosstalk between skin commensals and host immune cells *in vivo* and whether enhanced propionate export by *S. epidermidis* is a consequence of glucose limitation and/or cysteine/methionine bioavailability (**Figure 4A**). Differing fluxes through metabolic pathways for energy generation and metabolite export in the same organism under *in vivo* versus *in vitro* conditions can aid in designing strategies to coax microbes to export desirable metabolites such as SCFAs. This also highlights the need for model systems that better recapitulate the cutaneous microenvironment^30^ when investigating mechanisms by which commensals such as *S. epidermidis* contribute to skin health and disease.

We are the first study to generate *in vivo* gene expression data across a range of microbial species, individuals and skin sites to identify key antimicrobial functions and interactions. Although there are reports of skin commensals secreting antimicrobial compounds such as peptides or enzymes which can inhibit pathogens like *Staphylococcus aureus*, these studies were mostly conducted with *in vitro* or *ex vivo* models^7,8^. We confirmed that many of these products, such as the thiopeptide cutimycin^2^ and peptides of the lacticin 481 family^82^ which were described in previous studies, were also expressed *in vivo* on some individuals by *Cutibacterium acnes* and commensal *Staphylococcus* species respectively. Importantly, our dataset shows that the skin microbiome is a rich source of additional antimicrobial proteins/peptides whose activities and specificities remain uncharacterized. One intriguing example is a class of proteins harbouring a C terminal cysteine-rich region that is homologous to a class of bacteriocins known as the halocins. This class of bacteriocins have largely been characterized only in archaea but hypothetical proteins harbouring homologous regions have been found in the genomes of many *Staphylococcus* strains (https://www.ncbi.nlm.nih.gov/Structure/cdd/TIGR04449). Our data indicates that genes encoding these proteins with halocin C8-like domains are expressed by commensal *Staphylococcus* species *in vivo* (toe webs), providing a basis for further experimental validation of their antimicrobial properties. Since our workflow captures the expression of both host and microbial genes/pathways, correlating those with microbial DNA abundances and integrating these results with other functional annotations is a promising approach to identifying novel candidates mediating microbe-microbe and host-microbe interactions *in vivo*. We identified the *Malassezia restricta* gene DNF11_2196 as one such candidate owing to the strong negative correlation between transcript levels and *C. acnes* abundance, its predicted entry into the secretory pathway and its structural similarity to known protease inhibitors (**Figure 5**). Further application of this approach to population-scale cohorts could be useful for robust identification of physiologically relevant interactions between skin microbes and with host cells.

In conclusion, we have developed a systematic workflow to sample and analyze skin metatranscriptomes and have shown its utility for capturing *in vivo* microbial activities distinct from just DNA abundances, as well as identifying interactions that can shape microbial communities and host responses. Our data serves as a baseline for healthy individuals to enable future comparisons with datasets for disease states and gain deeper insights into microbial as well as host pathways that could be leveraged for diagnostic and therapeutic applications.

## Methods

### Subject recruitment

The study was conducted in the Genome Institute of Singapore (GIS). All associated protocols for this study were approved by the Agency for Science, Technology and Research Institutional Review Board (A*STAR IRB reference number 2021-094) on September 8th, 2021, and renewed until September 7th, 2025. All subjects recruited in this study were of Singaporean nationality or permanent residents, aged 21-65 and reported no skin disease at the time of sampling. Five subjects were recruited for cross-sectional and three for longitudinal analysis in a pilot cohort, respectively. For longitudinal analysis, subjects were sampled for 3 visits over 3 consecutive days with a minimum interval of 24 hours. For cross-sectional analysis, 27 subjects were recruited for the full cohort. All subjects were required to abstain from showering for at least 12 hours before sampling. Human skin biopsies from A*STAR Skin Research Laboratories (A*SRL) of 6 and 8mm were obtained from healthy skin donors, with approval from the NHG domain specific review board (NHG DSRB 2017/00224, NHG DSRB 2018/00945) at National University Health System (NUHS) and NSC domain specific review board (NSC DSRB 2019/00806) at National Skin Centre.

### Optimization of RNA Extraction with pilot cohort

Different sample collection methods, bead tubes, purification methods and various combinations of each were tested and are detailed in **Supplementary Data 1**. The different sample collection modes tested were with FLOQSwabs® (Copan Diagnostics; Cat. No. 502CS01) and tape discs (Cuderm; D-Squame D100). For sample collection with tape discs, tape discs were disinfected with 70% Ethanol for at least 1 minute and dried. Tape discs were applied to skin sites, peeled off and reapplied for a total of 50 iterations. Tape discs were then inserted into bead tubes, with the sticky surface facing the inside of the tube. For sample collection with swabs, swabs were first wetted with 1× phosphate buffer saline (PBS). The moistened swab was rotated and rubbed with constant pressure in a zig-zag pattern. This procedure was also repeated at an angle of 90° to the first rub, for a total of 1 minute. Swab contents were either dislodged by vigorous stirring in 800µL or 1300µL of DNA/RNA Shield (Zymo Research; Cat. No. ZYR.R1100) or were used directly for extraction. RNA extraction with RNeasy PowerMicrobiome kit (Qiagen; Cat. No. 26000) was carried out according to manufacturer’s instructions. RNA extraction with the Quick DNA/RNA Microprep Kit (Zymo Research; Cat. No. D7005) or Norgen BioTek RNA/DNA Purification Micro Kit (Norgen BioTek; Cat. No. 48700) was carried out according to their respective manufacturer’s instructions. Skin swabs were dislodged into DNA/RNA shield and bead-beating was carried out with ZR BashingBead Lysis tubes (0.1mm & 0.5mm), with 3 cycles of 1 min at 6m/s with 5 min interval on ice on the FastPrep-24 (MP Biomedicals) homogenizer. Lysates were used as inputs into these kits.

Samples which were extracted with hot sodium dodecyl sulfate (SDS) and hot acid phenol-chloroform were processed as follows. Skin samples were dislodged from swabs into DNA/RNA shield (2× concentrate; Zymo Research; Cat. No. R1200) and suspensions were added to 0.5× volume of SDS lysis solution (2% SDS, 16mM EDTA) pre-heated to 100°C. These mixtures were incubated at 100°C for 5 min with periodic mixing. Lysates were added to 1× volume of Acid-Phenol:Chloroform:Isoamyl alcohol, pH 4.5 (125:24:1) which was pre-heated to 65°C (Invitrogen; Cat. No. AM9720). This was mixed by vortexing and further incubated at 65°C for 10 min on a thermomixer with constant shaking at 1400rpm. Samples were transferred to 2mL screw cap tubes containing 550mg of autoclaved and air-dried 0.5mm zirconia/silica beads (BioSpec Products Inc.; Cat. No. 11079105Z). Bead-beating was carried out with 3 cycles of 1 min at 6m/s with 5 min interval on ice, using the FastPrep-24 homogenizer (MP Biomedicals). Samples were centrifuged for 5 min at 16,000g at 4°C and the aqueous phase was transferred to a new tube. 0.1× volume of 3M sodium acetate (pH5.5) (Invitrogen; Cat. No. AM9740), 2× volume of 100% ethanol and 1µL of linear acrylamide (5mg/ml; Invitrogen; Cat. No. AM9520) were added to the aqueous phase. RNA was allowed to precipitate overnight at -20°C and then centrifuged at 16,000g at 4°C for 5 min to obtain the pellet. RNA pellets were washed with 1mL of 70% ethanol and air dried for 5 min. Purified RNA was resuspended in RNase-free water, treated with an RNase-Free DNase Set (Qiagen; Cat. No. 79254) and cleaned up using RNeasy Min-Elute Clean up Kit (Qiagen; Cat. No. 74204).

Samples which were extracted with hot phenol were processed as follows. Skin samples were dislodged from swabs into 500µL DNA/RNA shield (Zymo Research; Cat. No. R1100). A volume of 800µL of phenol solution (Sigma Aldritch; Cat. No. P4682) pre-warmed to 65°C was added to the samples and gently mixed by pipetting without creating bubbles. The phenol-sample mixture was transferred to a tube containing 800µL of sodium acetate buffer solution (SAB; 50mM sodium acetate, 10mM EDTA, pH 5.2). The pre-lysate was transferred into a 2mL screw cap tube containing 550mg of 0.5mm zirconia/silica beads (BioSpec Products, Inc.; Cat. No. 11079105Z). Bead-beating on the FastPrep-24 (MP Biomedicals) homogenizer was carried out at 3 cycles of 1 min at 6.0m/s with 5 min intervals on ice. Lysates were incubated at 65°C for 30 min on a thermomixer and mixed at 1400rpm for 10 s every 1 min. Samples were centrifuged at maximum speed for 10 min at 4°C. The aqueous layer was added to 750µL of pre-warmed (65°C) phenol and incubated at 65°C for 5 min on a thermomixer, with mixing at 1400rpm for 10 s every 1 min. Samples were centrifuged at 16,000g for 10 min at 4°C and the aqueous layer was transferred to a tube containing 750µL of Acid-Phenol:Chloroform:Isoamyl alcohol, pH 4.5 (with IAA, 125:24:1) which was prewarmed to 65°C (Invitrogen; Cat. No. AM9720). The mixture was vortexed for 30 s and left at room temperature for 1 min. Samples were centrifuged at maximum speed for 10 min at 4°C. The aqueous layer was transferred to a new tube and 70µL of 3M sodium acetate and 700µL of isopropanol was added. The mixture was incubated at room temperature for 30 min and centrifuged at 16,000g for 30 min at 4°C to pellet the RNA. The supernatant was removed, and the RNA pellet was washed with 70% cold ethanol. Samples were centrifuged at 16,000g for 5 min, the ethanol was aspirated off and the pellet was air dried. The RNA pellet was then dissolved in 20µL of nuclease free water, treated with RNase-Free DNase Set (Qiagen; Cat. No. 79254) and cleaned up using RNeasy Min-Elute Clean up Kit (Qiagen; Cat. No. 74204).

Samples were extracted with the Direct-zol™ RNA MicroPrep kit (Zymo Research, Cat. No. ZYR.R2063) with Lysing matrix E (MP Biomedicals; Cat. No. 116914500) or ZR BashingBead Lysis tubes (0.1mm & 0.5mm; Zymo Research; Cat. No. S6012) or zirconia/silica beads (BioSpec Products; Cat. No. 11079105Z). Tape discs were directly added into bead tubes, while skin swabs or bacterial cells were dislodged into DNA/RNA shield. RNA purification was carried out according to the manufacturer’s protocol, except for some modifications such as the addition of 0.05% Tween 20 and using either 3 or 4 cycles of bead-beating, as detailed in **Supplementary Data 1**. Purified RNA was treated with RNase-Free DNase Set (Qiagen; Cat. No. 79254) and cleaned up using the RNeasy Min-Elute Clean up Kit (Qiagen; Cat. No. 74204).

### Sample collection for full cohort

Skin samples were collected using FLOQSwabs® (Copan Diagnostics; Cat. No. 502CS01) from five different skin sites (scalp, cheek, antecubital fossa, volar forearm, and toe web) from each subject (n=135 metagenomes and n=135 metatranscriptomes). For each skin site, three swabs were collected from the left and right side of the body and combined in a tube, except for the scalp, where only three swabs were used in total. Each swab was submerged in 1× phosphate buffer saline (PBS), and the excess solution was removed by pressing the swab against the tube wall. For each skin site, the moistened swab was rotated and rubbed, with constant pressure applied, in a zig-zag pattern and was repeated at an angle of 90° to the first rub, for a total of 1 min. The contents of the swab were dislodged by stirring vigorously in either 800µL or 1300µL of DNA/RNA Shield (Zymo Research; Cat. No. ZYR.R1100) for scalp and other skin sites respectively. Swabs were submerged in DNA/RNA shield for 5 min at room temperature and stirred vigorously again. Excess solution on each swab was collected by pressing the swab against the wall of the tube. This was repeated for the remaining swabs for each site. Negative controls (n=7) for each batch were collected by dislodging three swabs in DNA/RNA Shield (Zymo Research; Cat. No. ZYR.R1100) without sampling the skin. Each sample or control was split into 2 portions for RNA (approximately 600µL) and DNA (approximately 200µL) extraction. All samples were stored at -80°C prior to nucleic acid extraction.

### RNA extraction for metatranscriptomics (Direct-zol method)

RNA was extracted using the Direct-zol™ RNA MicroPrep kit (Zymo Research; Cat. No. ZYR.R2063). Compared to other extraction methods and kits, this approach was found to be the best performing in terms of RNA yield and RNA integrity (**Supplementary Data 1**). Bead-beating of samples in TRI Reagent® (Zymo Research; Cat. No. R2050-1-200) was done using ZR BashingBead Lysis Tubes (0.5 & 0.1mm; Zymo Research; Cat. No. S6012) and Fastprep-24 Instrument (MP Biomedicals) at 6.0m/s for a total of 3 min, in 1 min intervals, with 5 min incubation on ice between each interval. Samples were DNase treated and purified according to kit instructions. An additional DNase treatment was carried out by adding 2.5µL of DNase I and 10µL of RDD buffer from RNase-Free DNase Set (Qiagen; Cat. No. 79254), and 1µL of Recombinant RNasin RNase Inhibitor (10,000U; Promega; Cat. No. N2515) in a total volume of 100µL. The mixture was incubated at 37°C for 30 min and purified with RNeasy MinElute Cleanup Kit (Qiagen; Cat. No. 74204) in an elution volume of 14µL of RNase-free water. High Sensitivity RNA ScreenTape analysis (Agilent Technologies; Cat. No. 5067-5579, 5067-5580) was used to assess the quality of RNA and extracted RNAs were stored at -80°C.

### Mock communities

For metagenomic spike-ins, a mock community of 3 different bacteria (*Vibrio vulnificus* ATCC® 29307™, *Plesiomonas shigelloides* ATCC® 51903™, and *Listeria monocytogenes* ATCC® 35152™) was created by culturing each bacterial species in their respective growth media as recommended by ATCC. A growth curve was established for each bacterial species and the optical density (OD) of each culture was used to estimate the number of colony forming units (CFUs). Equal number of CFUs for each species were resuspended in DNA/RNA shield (Zymo Research; Cat. No. ZYR.R1100) and pooled together to create a mock community stock of concentration 1×10^5^ CFU/µL. Mock community stocks were aliquoted and stored at -80°C. Other mock communities for quality control testing were prepared to consist of 10 different bacterial species commonly part of human skin and gut microbiomes (**Supplementary Table 2**). Each species was cultured according to the recommended growth conditions and was counted using a hemocytometer. Cultures were centrifuged and each cell pellet was re-suspended in 100-500µL of DNA/RNA Shield (Zymo Research; R1100). Samples were stored at -80°C prior to combining the cells according to the abundances summarized in **Supplementary Table 2**.

### DNA extraction for metagenomics (EZ1 method)

This approach (EZ1 method) was used for DNA extraction due to relatively poor DNA yields from the Direct-zol method, which is optimized for RNA extraction. Spike-ins of the metagenomic mock community (1.5×10^4^ CFUs) were introduced to each sample prior to DNA extraction. Lysis of samples was carried out by adding 500µL of ATL Buffer (Qiagen; Cat. No. 19076) to the sample and homogenisation in Lysing Matrix E tubes (MP Biomedicals; Cat. No. 116914500) with a FastPrep-24 Instrument at a speed of 6.0m/s for 40 s, done twice in total. Cell debris was pelleted at 16,000g for 5 min and the supernatant was treated with 12µL of Proteinase K at 56°C for 15 min prior to purification with EZ1 DNA Tissue Kit (Qiagen; Cat. No. 953034) using the EZ1 Advanced XL machine (Qiagen). A Qubit fluorimeter was used to quantify the amount of DNA.

### Extraction quality control

To assess the lysis efficiencies of different protocols, each mock community (**Supplementary Table 2**) was extracted with EZ1 DNA Tissue Kit (Qiagen; 953034; EZ1 method) and a separate aliquot with TRI Reagent® (Zymo Research; Cat. No. R2050-1-200; Direct-zol method). DNA was obtained from the Direct-zol method using additional steps after lysis according to the TRIzol™ Reagent (DNA isolation) User Guide. DNA was cleaned up and concentrated with 2× AMPure XP beads (Beckman Coulter; A63882). DNA was eluted in 30µL of Buffer EB (Qiagen; 19086). Qubit RNA High Sensitivity assay kit (Thermo Fisher Scientific; Q32852) was used to check for RNA contamination. Samples extracted by the Direct-Zol method that had detectable levels of RNA were treated with RNase A (100mg/mL; Qiagen; 19101), cleaned up with 2× AMPure XP beads, and eluted in 30µL of Buffer EB. DNA was used for library preparation and sequenced on a Illumina Novaseq X (2×150 bp reads). This data was used to confirm that metagenomic relative abundances obtained based on DNA extracted from both methods were highly consistent across the different mock communities that were tested (Pearson’s *R*=0.95; **Supplementary** Figure 23).

### Preparation of ribosomal RNA depletion mix

Five different oligo probe pools (desalted) were ordered from Integrated DNA Technologies (IDT) targeting either *Malassezia* rRNA or other fungal rRNAs (**Supplementary Table 1**). The probe pools for *Malassezia* rRNAs were mixed to form a pool of 868 probes, while probe pools for other fungal rRNAs formed a pool of 630 probes. Both oligo pools were resuspended to a concentration of 2 µM/probe. The two oligo probe pools were then mixed together (i.e., *Malassezia*:other fungi – 4:1) to form a pan-fungal rRNA depletion probe pool. The final rRNA depletion probe pool was made by combining the NEBNext rRNA Depletion Solution from NEBNext rRNA Depletion Kit V2 (Human/Mouse/Rat; New England Biolabs; Cat. No. E7405) and NEBNext rRNA Depletion Kit V2 (Bacteria; New England Biolabs; Cat. No. E7850) with the custom pan-fungal rRNA depletion probe pool at a volume ratio of 40:9:1, with 2 µL of this final probe mix used per sample for rRNA depletion.

### RNA library preparation

Human and microbial ribosomal RNAs were depleted from 5–10 ng of total RNA or the entire volume of eluted total RNA. Libraries were prepared according to manufacturer’s instructions using NEBNext Ultra II Directional RNA Library Prep Kit for Illumina (New England Biolabs; Cat. No. E7760). Depending on the RIN value of RNA, either Section 2 or 3 of the protocol was used. Library enrichment was carried out using NEBNext Multiplex Oligos for Illumina (96 Unique Dual Index Primer Pairs; New England Biolabs; Cat. No. E6440) or NEBNext Multiplex Oligos for Illumina (Unique Dual Index UMI Adaptors RNA Set 1; New England Biolabs; Cat. No. E7416) with 14 cycles of enrichment PCR. The quality of libraries was assessed by High Sensitivity D1000 ScreenTape Assay (Agilent Technologies; Cat. No. 5067-558). Libraries were pooled in equimolar proportions and sequenced on a Illumina HiSeq X Ten system (∼35 million 2×150bp read pairs per library).

### DNA library preparation

NEBNext Ultra II FS DNA Library Prep Kit for Illumina (New England Biolabs; Cat. No. E7805) was used according to manufacturer’s instructions with some modifications. A volume of 26µL of DNA was used as input and subjected to 10 min of fragmentation at 37°C. Fragmented DNA was used for adapter ligation and was cleaned up using 0.6× volume of AMPure XP Reagent (Beckman Coulter; Cat. No. A63882). Adaptor ligated DNA was amplified for 12 cycles using NEBNext Multiplex Oligos for Illumina (96 Unique Dual Index Primer Pairs; New England Biolabs; Cat. No. E6440) and cleaned up with 0.7× volume of AMPure XP Reagent (Beckman Coulter; Cat. No. A63882). The final library was eluted in 20µL of EB Buffer (Qiagen; Cat. No. 19086). Quality of libraries were assessed by the High Sensitivity D1000 ScreenTape Assay (Agilent Technologies; Cat. No. 5067-558). Libraries were pooled in equimolar proportions and sequenced on a Illumina HiSeq X Ten system (∼25 million 2×150bp read pairs per library).

### Data pre-processing and quality control

Short reads from metagenomic and metatranscriptomic libraries were processed using a Nextflow^83^ pipeline (https://github.com/Chiamh/meta-omics-nf). Quality control and adapter trimming were done using fastp^84^ (v0.22.0) with default settings. Metagenomes were further pre-processed by mapping to the hg38 human reference genome using BWA-MEM^85^ (v0.7.10-r789) and reads that failed to map to hg38 were extracted using samtools^86^ (v1.13) with parameters −f12 −F256. Human RNA was removed from metatranscriptomes by mapping to hg38 using STAR^87^ (2.7.9a). Reads originating from microbial ribosomal RNAs (rRNAs) were computationally removed from metatranscriptomes using bbduk.sh (BBMap v38.93) and a k-mer database for rRNAs. Subsequently, microbial RNA reads were de-duplicated using clumpify.sh (BBMap v38.93) with parameters dedupe=t and optical=f.

### Taxonomic classification

Metagenomic reads were classified using Kraken2^24^ (v2.1.2) and Bracken^88^ (v2.6.1) with parameters --use-names, --paired and --report-minimizer-data. Metatranscriptomic reads were classified with Kraken2 using the same parameters. A 50Gbp Kraken2 database built from RefSeq bacterial, archaeal, viral, fungal and human (hg38) genomes, as well as plasmid sequences was used. This database also contains additional *Malassezia* assemblies downloaded from NCBI (**Supplementary Data 11**). Samples with at least 10,000 paired reads were retained and reads that were still taxonomically assigned to *Homo sapiens* were removed. Microbial reads were defined as the sum of reads classified as bacteria (taxid 2), archaea (taxid 2157), virus (taxid 10239), and fungi (taxid 4751).

False positive species assignments for metagenomic reads were identified and removed using an approach similar to Breitwieser et al^23^. Species assignment of metagenomic reads were considered true positives if there were ≥2000 unique Kraken2 minimizers per 1 million microbial reads or had ≥10 read pairs for the species with ≥10× more unique minimizers than read pairs. False positive species assignments for metatranscriptomic reads were identified and removed using a similar approach, with empirically determined minimizer thresholds. Specifically, RNAs from three biological repeats (MHS445, MHS589 and MHS590), each comprising a mix of non-skin bacteria *Plesiomonas shigelloides*, *Vibrio vulnificus* and *Listeria monocytogenes* were extracted, processed, and sequenced in the same way as other metatranscriptomes in this study. Random subsamples of these libraries were added to a skin metatranscriptome (MHS413) such that read pairs from *P. shigelloides*, *V. vulnificus* and *L. monocytogenes* were each present at relative abundances of approximately 0.1%, 1% or 10% of all reads in the metatranscriptome. These metatranscriptome mixtures were subsequently rarefied to either 10^4^ or 10^6^ read pairs and reads were classified using Kraken2. For each true positive spike-in species (*P. shigelloides, V. vulnificus and L. monocytogenes*), false positive classifications of reads to other species belonging to the genera *Listeria*, *Plesiomonas*, *Vibrio*, *Pseudomonas*, *Salmonella* and *Klebsiella* were identified. The log_10_ ratio of unique species-specific minimizers per million microbial reads was plotted against log_10_ paired counts of each species to determine that 10^4^ unique minimizers per million microbial reads could distinguish most true positive species assignments from false positives (**Supplementary** Figure 2C). Species assignment of metatranscriptomic reads were considered true positives if there were ≥10^4^ unique Kraken2 minimizers per 1 million microbial reads or had ≥10 read pairs for the species with ≥10× more unique minimizers than read pairs. Species contributing <0.1% RNA relative abundance were removed from metatranscriptomes to avoid false positives from incorrect read classification. Species contributing <0.1% DNA relative abundance were also removed from each metagenome unless they were also present in the matched metatranscriptome (**Supplementary** Figure 2C).

### Kitome removal

Potential reagent and laboratory contamination-associated species (the “kitome”) were identified and removed via a multi-step process (**Supplementary** Figure 2A). Swab extraction controls (n=7 negative controls) were sequenced from fresh swabs unexposed to human skin. Species with ≥0.1% relative abundance in any negative control constituted an initial list of 70 candidate genera. Each of these 70 genera were classified as kitome candidates if they were not previously reported on skin or in skin diseases, based on hits to the Disbiome database^89^, the MicrophenoDB^90^ or a PubMed literature search (last accessed 24^th^ Nov 2022) using the search terms ([Genus name] AND "Skin" AND "microbiome" AND "human"). Pairwise Pearson correlations between all species in the metagenomes or metatranscriptomes and these kitome candidates were calculated using SparCC^91^. Any species with a strong positive correlation (*r*≥0.8) with kitome candidate species were also marked as potential kitome contaminants and removed from downstream analyses. We observed that abundances of previously reported^92^ environmental and kit contaminants such as species of the genera *Achromobacter, Bradyrhizobium, Mycolibacterium, Mycobacterium* and *Brevundimonas* were not strongly positively correlated with skin or oral microbes and could hence be easily distinguished from them for removal (**Supplementary** Figure 2B).

### Functional classification

Metagenomic and metatranscriptomic reads were functionally classified based on a similar strategy to that of HUMAnN3^20^, using a custom Nextflow pipeline (https://github.com/Chiamh/meta-omics-nf). Reads were first aligned in single-end mode using Bowtie2^93^ (v2.4.4) in –very-sensitive mode, to the Integrated Human Skin Microbial Gene Catalog^19^ (IHSMGC) comprising approximately 10.9 million non-redundant genes. A coverage filter of 50% across the length of any given hit pangene sequence was used^20^. Reads which failed to align to the IHSMGC were mapped against the UniRef90^94^ database (downloaded 9^th^ June 2021) using Diamond^95^ (v2.0.12) with parameters blastx, --id 80 --query-cover 90 and --max-target-seqs 1. For searches against the UniRef90 database, only alignments with ≥80% sequence identity, ≥90% query (read) coverage and ≥50% subject (UniRef90 representative sequence) coverage were considered as hits. These thresholds were the same as the defaults used by HUMAnN3 and were previously shown to increase specificity of alignments to the UniRef90 database with a relatively small reduction in sensitivity^20^.

Pangenes and UniRef90 clusters with valid hits after mapping were annotated and grouped into orthologous groups (OGs) using EggNOG mapper^96^ (v2.1.6) and the EggNOG 5.0 database^97^ with parameters -m diamond and --go_evidence all. This provides annotations for Gene Ontology (GO)^98,99^ and Kyoto Encyclopaedia of Genes and Genomes^100^ (KEGG) pathway analysis. Gene-level analysis was done for OGs by summarizing the read counts at the level of bacteria (taxid 2) or fungi (taxid 4751). Rarefaction analysis for bacterial and fungal OGs was conducted in R using the ‘rarecurve’ function from the vegan package (v2.6-6.1).

### Pathway abundance and contributional diversity analysis

Pathway abundances were computed using HUMAnN3 (v3.8), using a custom structured KEGG module definition file (https://github.com/CSB5/skin_metatranscriptome), with each definition retrieved using the KEGG REST API (e.g. https://rest.kegg.jp/get/M00357). Microbial alpha (Simpson) and beta (Bray-Curtis dissimilarity) diversity for each KEGG module was calculated as described in Franzosa et al^9^. Only modules that were core to a skin site (non-zero counts in >75% of individuals) with largely known microbial provenance (<25% unclassified species-level reads per module) were used for this analysis. Alpha or beta diversity scores >0.5 were considered “high” diversity.

### Staphyloccocus epidermidis culture experiments

Overnight cultures of *S. epidermidis* (ATCC 12228) were diluted into 3 volumetric flasks of culture at OD_600_ 0.01 and incubated at 37°C for 6h to log phase (OD_600_ 0.4-0.5). Each flask was split into 3 tubes (biological triplicates) and subjected to different stress exposures as described in Avican et al^42^ (**Supplementary Table 2**). A portion (2×500µL) of each culture was transferred into 2mL tubes after stress exposure and 1.5mL of DNA/RNA Shield (Zymo Research; Cat. No. R1100) was added into each tube and mixed well. Cell mixtures were then centrifuged at 8000g for 10min at 4°C to pellet cells. Each cell pellet was re-suspended in 500µL of TRI Reagent (Zymo Research; Cat. No. R2050-1-200) and each technical duplicate was combined into 1 tube with a final volume of 1mL. Samples were stored at -80°C prior to RNA extraction.

### Alignment to species-specific pangenomes

Bacterial species-specific analyses was done by mapping metagenomic and metatranscriptomic reads to curated pangenomes^101^ (https://ngdc.cncb.ac.cn/propan/) of eight commonly found skin microbes (*Staphylococcus aureus, Staphylococcus epidermidis, Staphylococcus hominis, Staphyloccocus capitis, Cutibacterium acnes, Cutibacterium modestum, Corynebacterium tuberculostearicum and Corynebacterium ureicelerivorans*), together with decoy genomes of the non-skin microbes *Achromobacter xylosoxidans, Plesiomonas shigelloides, Vibrio vulnificus* and *Listeria monocytogenes*. All genes in the eight skin species pangenomes were further clustered using CD-HIT (v4.8.1) at ≥95% protein sequence identity and mutual alignment coverage ≥90%, following guidelines by Li et al^19^. Reads were aligned to the de-replicated eight skin species pangenomes in single-end mode using Bowtie2 (v2.4.4) in --very-sensitive mode (similar to HUMAnN3). A coverage filter of 50% across the length of any given hit pangene sequence was used. Fungal species-specific analyses were done by pseudo-alignment of metatranscriptomic reads to the reference transcriptomes of multiple *Malassezia* species, with their reference genomes as decoys to minimize occurrences of non-transcriptomic reads being erroneously counted due to similarities to the annotated transcriptome, following recommendations for Salmon^102^ (v1.10.1). Unlike fungal reads, bacterial reads were mapped in single-end mode to account for the organization of ORFs in polycistronic mRNAs, similar to the approach adopted by HUMAnN3. Single-ended read coverage over bacterial coding sequences was computed using picard (v3.1.1) CollectRnaSeqMetrics with arguments -STRAND FIRST_READ_TRANSCRIPTION_STRAND. Paired-end read coverage over fungal transcripts were computed using picard CollectRnaSeqMetrics with arguments - STRAND SECOND_READ_TRANSCRIPTION_STRAND.

### Transcriptional activity analysis

An organism’s transcriptional activity was calculated in a similar way as described in Abu-Ali et al^13^. Metagenomic or metatranscriptomic reads were mapped to the IHSMGC^19^ pangene catalogue and the UniRef90 database. Read counts for any given species (including all sub-species) were then divided by the length of each feature in kilobases to obtain the number of reads per kilobase (RPK). Reads belonging to a species that could not be mapped to pangene or UniRef90 features were assumed to belong to an unknown gene of length 1 kilobase for computing RPK. RPKs for each metagenomic or metatranscriptomic library were then summed and divided by 10^6^ to obtain a per-sample scaling factor. Metatranscriptomic transcripts per million (TPM) or metagenomic copies per million (CPM) values were computed by dividing a feature’s RPK with the per-sample scaling factor. The transcriptional activity of a species was estimated by summing species-level TPM values (RNA) and dividing them by species-level CPM (DNA) values.

### Differential expression analysis

Differential expression analysis was done using DESeq2^103^ (v1.36.0). Raw counts were summarized at the level of bacterial (taxid 2) and fungal (taxid 4751) orthologous groups for gene-level differential expression analysis. The design formula was ‘∼ subject ID + skin site (Sc, Ch, Ac, Vf or Tw) + assay (RNA or DNA) + skin site:assay’. This design formula accounts for within-subject dependencies and variations in gene copy numbers in metagenomes while testing for differences in microbial gene expression between skin sites. Only features (rows) with median read count ≥10 for both DNA and RNA were kept. Size factors were estimated separately for the metagenomic and metatranscriptomic count matrices using the ‘poscounts’ function to account for data sparsity. Differential expression analysis at individual species-level between two *in vivo* conditions was similarly done, except that only metatranscriptomic read counts were used from libraries with ≥200,000 species-specific reads, and the design formula was ‘∼ subject + skin site’. Differential expression analysis for *Staphylococcus epidermidis* comparing *in vivo* versus *in vitro* growth conditions was similarly done, except that the design formula was ‘∼ batch + growth condition’ to account for experimental batch effects when using *in vitro* RNA-seq data from different sources. Batch corrected principal component analysis (PCA) plots were derived from inputs processed with the removeBatchEffect function from limma^104^ (v3.60.4).

### Gene set enrichment analysis

Gene set enrichment analysis (GSEA) for differentially expressed microbial features was done using clusterProfiler^105^ (v4.4.4) with the arguments: eps=0, nPermSimple=10000 and seed=TRUE.

### Integration of metatranscriptomics data with metabolic models

Genome-scale metabolic models (GSMMs) for *Staphylococcus epidermidis* ATCC 12228 and *Propionibacterium acnes* KPA171202 were obtained from the AGORA database^110^. These models were constrained using COBRApy^111^ based on specific exchange flux values corresponding to the conditions under which the simulations were performed. Genes in the metabolic models were mapped based on transcript levels, calculated as the geometric mean of transcript abundance (TPMs) across replicates or samples under the same growth condition. The integration of these TPM values into genome-scale models and the subsequent flux balance analysis (FBA) was carried out in Python (v3.12) using RIPTiDe^112^ (v3.4.81). Briefly, RIPTiDe incorporates gene expression data using reaction parsimony, generating context-specific GSMMs. The context-specific models were simulated using FBA to identify flux distributions within the organism under each condition separately. Non-metric multidimensional scaling (NMDS) of the Bray-Curtis distances between flux distributions was performed using functions from the vegan package (v2.6-4) in R (v4.3.0) to compare the flux profiles across conditions. Differentially abundant reactions were identified using a generalized linear model (GLM: reactions ∼ group), with reactions showing adjusted p-values below 0.001 and absolute estimate values greater than 2 being considered significantly different between conditions. These reactions were plotted using ggplot2 (v3.5.1) in R.

### Identification and analysis of antimicrobial genes

Hidden Markov Models (HMMs) of various classes of antimicrobial proteins were downloaded from NCBI (https://www.ncbi.nlm.nih.gov/protfam; **Supplementary Data 12**). Microbial pangenes and representative sequences from UniRef90 gene clusters were searched against these HMMs using hmmscan from hmmer^106^ (v3.3.2). Hits to a HMM were only kept if both the “seq” and “best one domain” scores were greater than or equal to the sequence and domain cutoffs given by NCBI. Antimicrobial genes were considered “present” in metatranscriptomes if their read counts were ≥5, with ≥50% coverage over the gene body. Multiple sequence alignment (MSA) of microbial sequences and representative sequences from HMMs was done using MUSCLE^107^ (v5.1). Matrices of pairwise percentage identities were computed from the MSAs and coverage statistics were computed using a custom python script (https://github.com/CSB5/skin_metatranscriptome).

### Identification of microbe-gene correlations

SignalP^108^ (v6.0) was used to classify microbial proteins (features) that can enter the secretory pathway. Fungal and bacterial proteins were analysed in “fast” mode with the options --organism “eukarya” or “other” respectively. Microbial features predicted to enter the secretory pathway were shortlisted for correlation analysis. For a given pair of microbes at a skin site, pairwise Spearman correlations were computed between the variance stabilized counts (RNA) of microbial features computed from DESeq2 and the centered log-ratio^109^ (clr) transformed counts (averaged across 1000 Monte Carlo instances) of microbial abundances (DNA) computed from aldex2 (v1.28.1) and the aldex2propr function from propr (v2.1.2). Features were considered significantly correlated only with Spearman’s ρ≥0.7 and FDR adjusted p-value≤0.05.

### Structural similarity searches

Protein Data Bank (PDB) files of selected microbial proteins were downloaded from the AlphaFold Protein Structure Database (https://alphafold.ebi.ac.uk/). Structural similarity searches were done using the Foldseek Search server (https://search.foldseek.com/search) in 3Di/AA mode and the Dali server (http://ekhidna2.biocenter.helsinki.fi/dali/) in PDB search mode.

### Microbial strain isolation

Commensal *Staphylococcal* strains were isolated from skin swabs for 4 healthy donors across 4 separate body sites: antecubital fossa, axilla, cheek and scalp. Swabs were immersed in 2.5mL BHI broth (Oxoid), incubated at 37°C at 210rpm for 2 hours before being spread onto TSA-SB (Thermo Scientific) and Baird Parker (Oxoid) plates and incubated at 37°C for 24 hours. Colonies were then picked and grown in 4ml BHI broth for 16 hours overnight at 230rpm and 37°C. 200μL of the cultures were spun down at 5000rpm for 5 minutes. To extract gDNA the pellet was resuspended in 50μL QuickExtract (Lucigen). Bacterial suspensions were then heated to 65°C for 6 minutes, briefly vortexed, then heated to 98°C for 4 minutes and briefly vortexed again. PCR reactions were setup with 10μL KAPA SYBR FAST (Merck), 6.4μL nuclease free water (Promega), 3μL extracted gDNA and 0.6μL primer mix. Primer mixes were specific for 4 specific *Staphylococcal* species (**Supplementary Table 1**). PCR was run on an AriaMx Real Time PCR System (Agilent Technologies). PCR products were sent for Sanger sequencing to confirm product sequences. The identified *Staphylococcal* colonies were inoculated into 4ml BHI broth and grown overnight for 16 hours at 230rpm and 37°C. Optical density (OD) was measured with a SpectraMax M5 Microplate Reader (Molecular Devices). Overnight cultures were then grown from OD 0.1 in fresh BHI for 26 hours at 230rpm and 37°C. Final OD measurements for all strains after 26 hours were normalized to OD 2.75. Bacterial cultures were spun down at 5000rpm for 5 minutes, pellets were discarded and supernatants stored at 4°C.

### Human keratinocyte cell culture

A N/TERT keratinocyte cell line was used for ELISA experiments. A genetically modified N/TERT keratinocyte cell line was utilized for Nano-Glo HiBiT experiments which contained a 33 nucleotide HiBiT tag directly adjacent to the *IL1B* start codon. Keratinocytes were cultured in keratinocyte serum free medium (KSFM; Gibco) supplemented with bovine pituitary extract (BPE) at a final concentration of 20μg/mL, epidermal growth factor (EGF) at a final concentration of 0.2ng/mL, calcium chloride at a final concentration of 300μM and 1:1000 penicillin-streptomycin (Gibco). Keratinocytes were seeded onto 96 well cell culture plate (Greiner) at a density of 20,000 cells per well and grown for 24 hours in 100μL KSFM. After 24 hours KSFM was removed and the keratinocytes were cultured in 100μL KSFM with either 1μM anisomycin (Merck), 0.2% TritonX (Merck), 5% BHI or 5% *Staphylococcal* supernatants. Keratinocytes were cultured overnight for 16 hours in treatment conditions.

### Nano-Glo HiBiT for pro-IL-1B measurements

After 16 hours of treatment of the HiBiT N/TERT keratinocytes, 50μL of the KSFM was transferred to a white-bottom 96 well plate (Sigma-Aldrich) and mixed with 50μL Nano-Glo HiBiT extracellular reagent. Plates were mixed on an orbital shaker for 1 minute, incubated for 10 minutes at room temperature before luminescence was measured with a SpectraMax M5 Microplate Reader.

### ELISA for cleaved IL-1B

After 16 hours of treatment of the N/TERT keratinocytes, 100μL of the KSFM was transferred to a new 96 well cell culture plate and stored at -80 °C until required. Human IL-1B ELISA were performed in accordance with the manufacturer’s instructions (FineTest), OD measurements were performed with a Spark Multimode Microplate Reader (Tecan) at 450nm and corrected against 570nm.

### Statistical analysis and visualization

Statistical tests and visualizations were done using R (v4.2.0), ggplot (v3.3.6), EnhancedVolcano (v1.14.0) and ggpubr (v0.4.0).

## Data availability

Shotgun metagenomic and metatranscriptomic sequencing data is available from the European Nucleotide Archive (ENA – https://www.ebi.ac.uk/ena/browser/home) under project accession number PRJEB82796. All large datasets, Foldseek webserver outputs, Dali webserver outputs, the Kraken2 database used for taxonomic classification, and other gene annotation databases are available on Figshare at https://figshare.com/projects/Skin_metatranscriptomics_manuscript_2024/202683.

## Code availability

Source code for scripts used to analyze the data are available at https://github.com/CSB5/skin_metatranscriptome.

## Acknowledgements

The authors wish to thank Lim Thiam Chye and Nelson Teo from NUHS, and Steven Thng and Yew Yik Weng from NSC for collecting patient skin biopsies. This work is supported by the Asian Skin Microbiome Programme 2.0 (Industry Alignment Fund Pre-Positioning; H22J1a0040), Agency for Science, Technology and Research BMRC EDB IAF-PP grants – H17/01/a0/004 and Agency for Science, Technology and Research BMRC Central Research Funds (ATR) and a National Medical Research Council (NMRC) Clinician Scientist-Individual Research Grant (CIRG23jul-0018). This work was also supported by the A*STAR Computational Resource Centre through the use of its high-performance computing facilities. The N/TERT keratinocyte cell lines were provided by the Zhong Lab, from LKC-Medicine to A*SRL.

