## Supplementary Figures for "Skin metatranscriptomics reveals landscape of variation in microbial activity and gene expression across the human body"

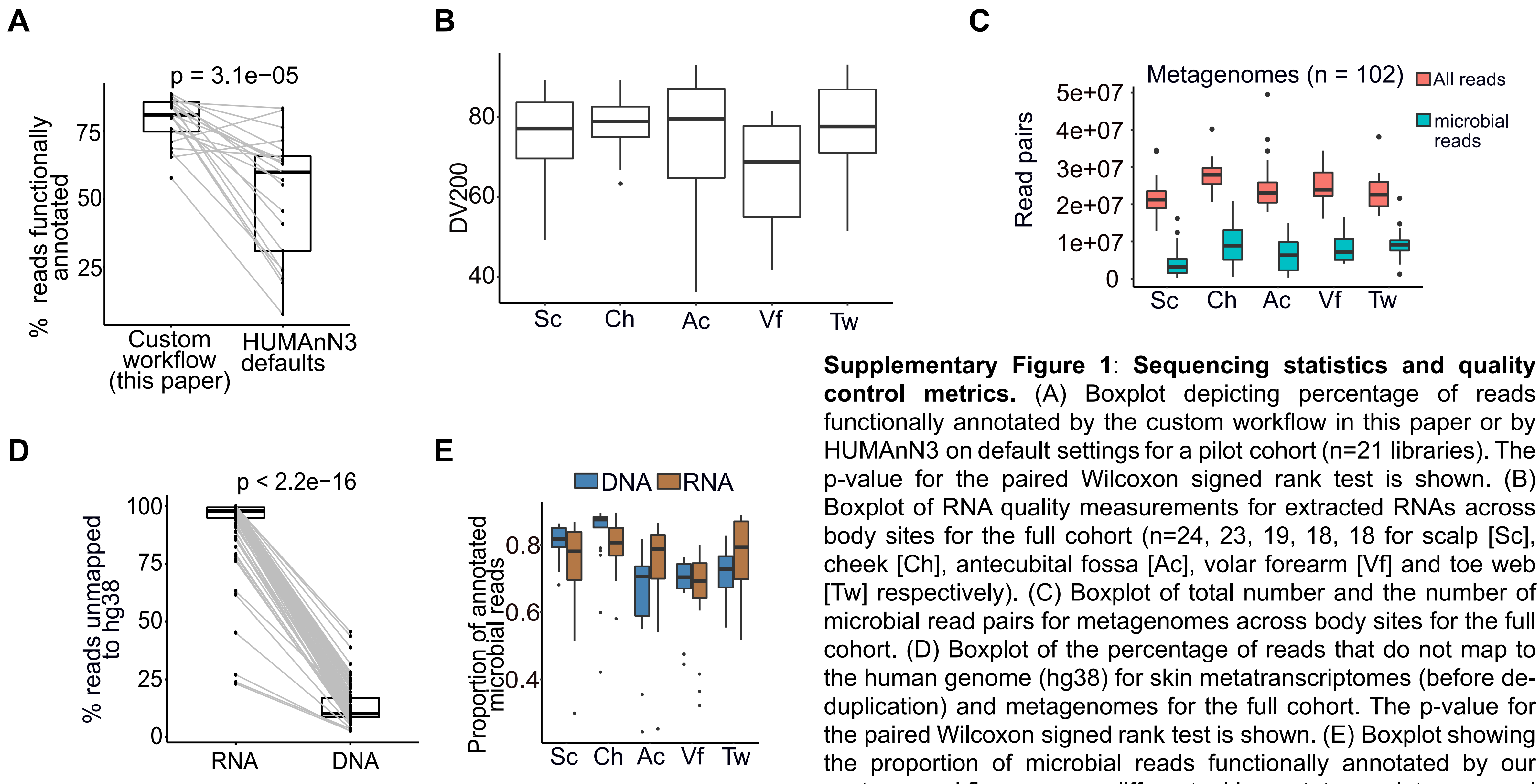


**Supplementary Figure 1**: **Sequencing statistics and quality control metrics.** (A) Boxplot depicting percentage of reads functionally annotated by the custom workflow in this paper or by HUMAnN3 on default settings for a pilot cohort (n=21 libraries). The p-value for the paired Wilcoxon signed rank test is shown. (B) Boxplot of RNA quality measurements for extracted RNAs across body sites for the full cohort (n=24, 23, 19, 18, 18 for scalp [Sc], cheek [Ch], antecubital fossa [Ac], volar forearm [Vf] and toe web [Tw] respectively). (C) Boxplot of total number and the number of microbial read pairs for metagenomes across body sites for the full cohort. (D) Boxplot of the percentage of reads that do not map to the human genome (hg38) for skin metatranscriptomes (before de-duplication) and metagenomes for the full cohort. The p-value for the paired Wilcoxon signed rank test is shown. (E) Boxplot showing the proportion of microbial reads functionally annotated by our custom workflow across different skin metatranscriptomes and metagenomes for the full cohort.

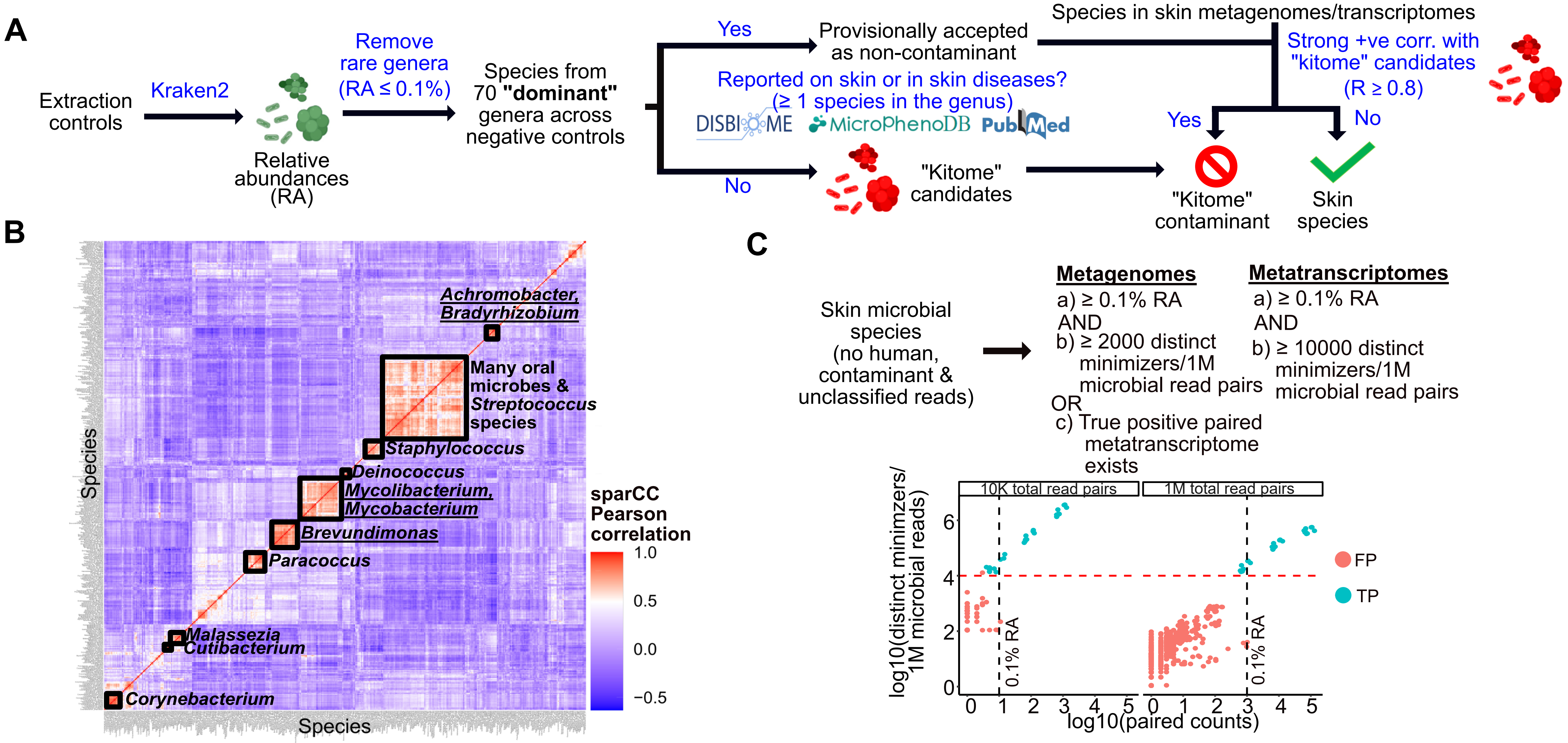


**Supplementary Figure 2: Controlling for experimental and computational artifacts in skin metatranscriptomes.** (A) Summary of steps for the identification of kit contaminants from negative control libraries. (B) Heatmap showing that the metagenomic abundances of contaminant genera (underlined) are not correlated with skin and oral microbes, facilitating their identification and removal. (C) Criteria for differentiating true positive (TP) from false positive (FP) species assignments from Kraken2 using thresholds for relative abundance (black dotted line) and for log10 distinct minimizers for a given species per 1M microbial reads (red dotted line). These thresholds are robust to library size (10K and 1M total read pairs).

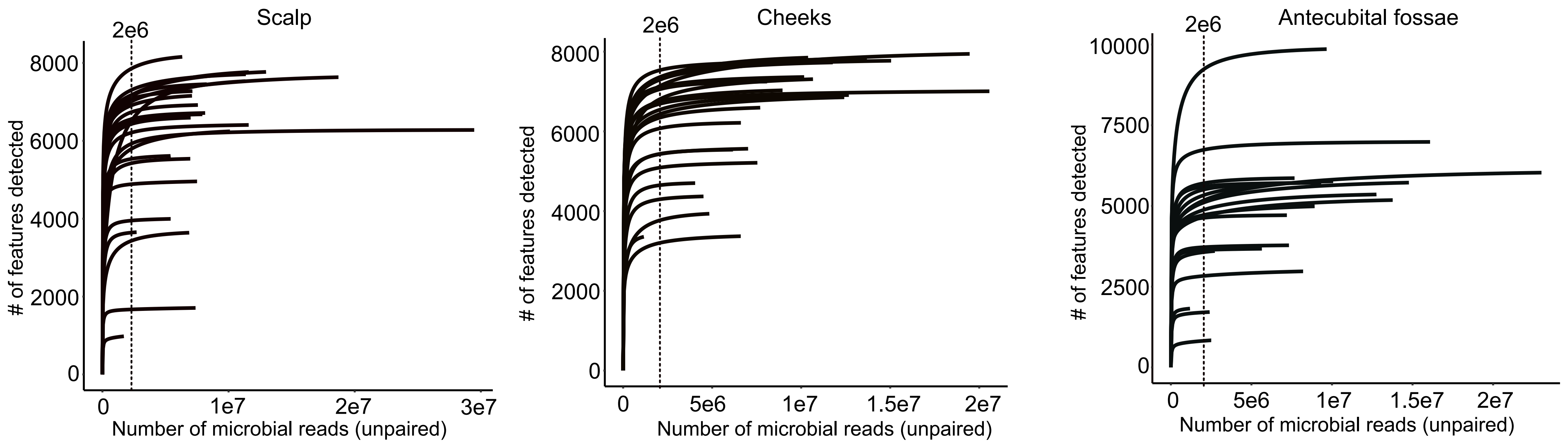


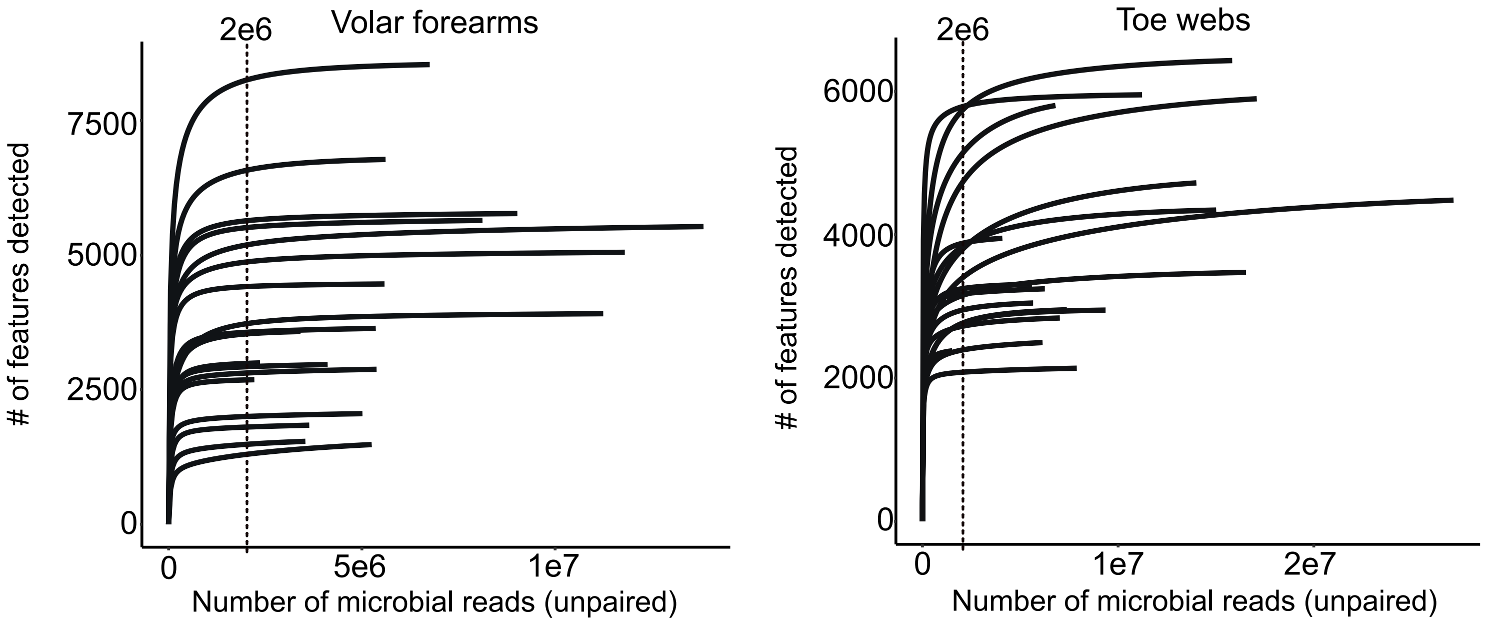


**Supplementary Figure 3: Rarefaction analysis of expressed microbial features.** Rarefaction curves of transcribed bacterial and fungal orthologous groups (OGs) for samples across the five different skin sites, as a function of the number of microbial reads sequenced. Most curves have plateaued, suggesting that most transcribed features were sequenced in the corresponding samples. Data is from the full cohort (n=24, 23, 19, 18, 18 for scalp, cheek, antecubital fossa, volar forearm and toe web respectively). A threshold of 2 unpaired million microbial reads (1 million read pairs) is shown.

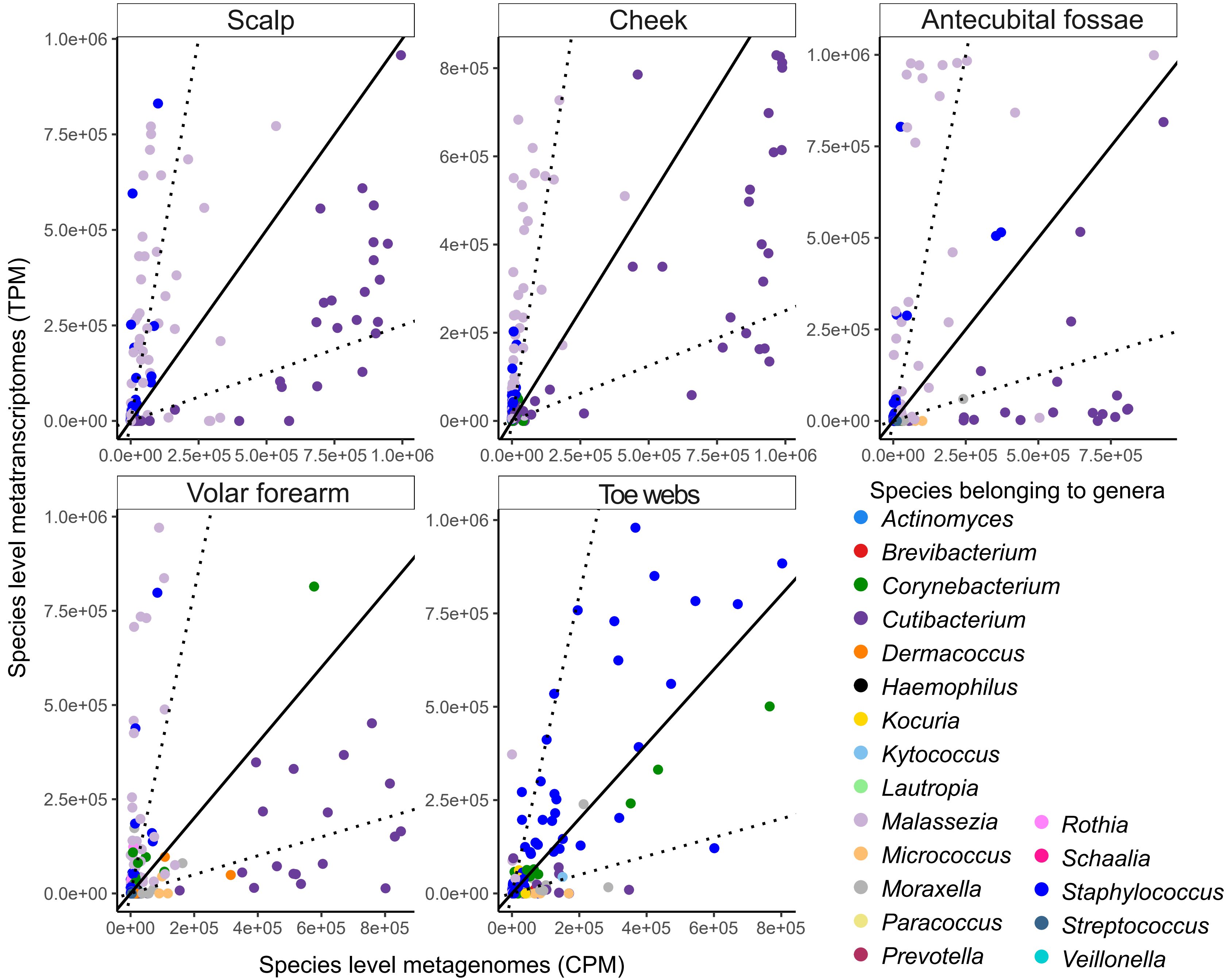


**Supplementary Figure 4: Comparison of metatranscriptomic and metagenomic counts for various species across skin sites.** Scatter plots of species level metatranscriptomic (RNA) counts expressed as transcripts per million (TPM) against species level metagenomic (DNA) counts expressed as counts per million (CPM). Points on the solid line represent equal proportions of RNA to DNA for a species. The areas beyond the dotted lines represents ≥4-fold differences between RNA and DNA counts. Plots represent data from all 102 paired metagenomes and metatranscriptomes in this study. Note that *Staphylococcus* (dark blue) *and Malassezia* (lilac) species have an outsized contribution to metatranscriptomes at most sites despite their lower representation in metagenomes. Data is from the full cohort (n=24, 23, 19, 18, 18 for scalp, cheek, antecubital fossa, volar forearm and toe web respectively).

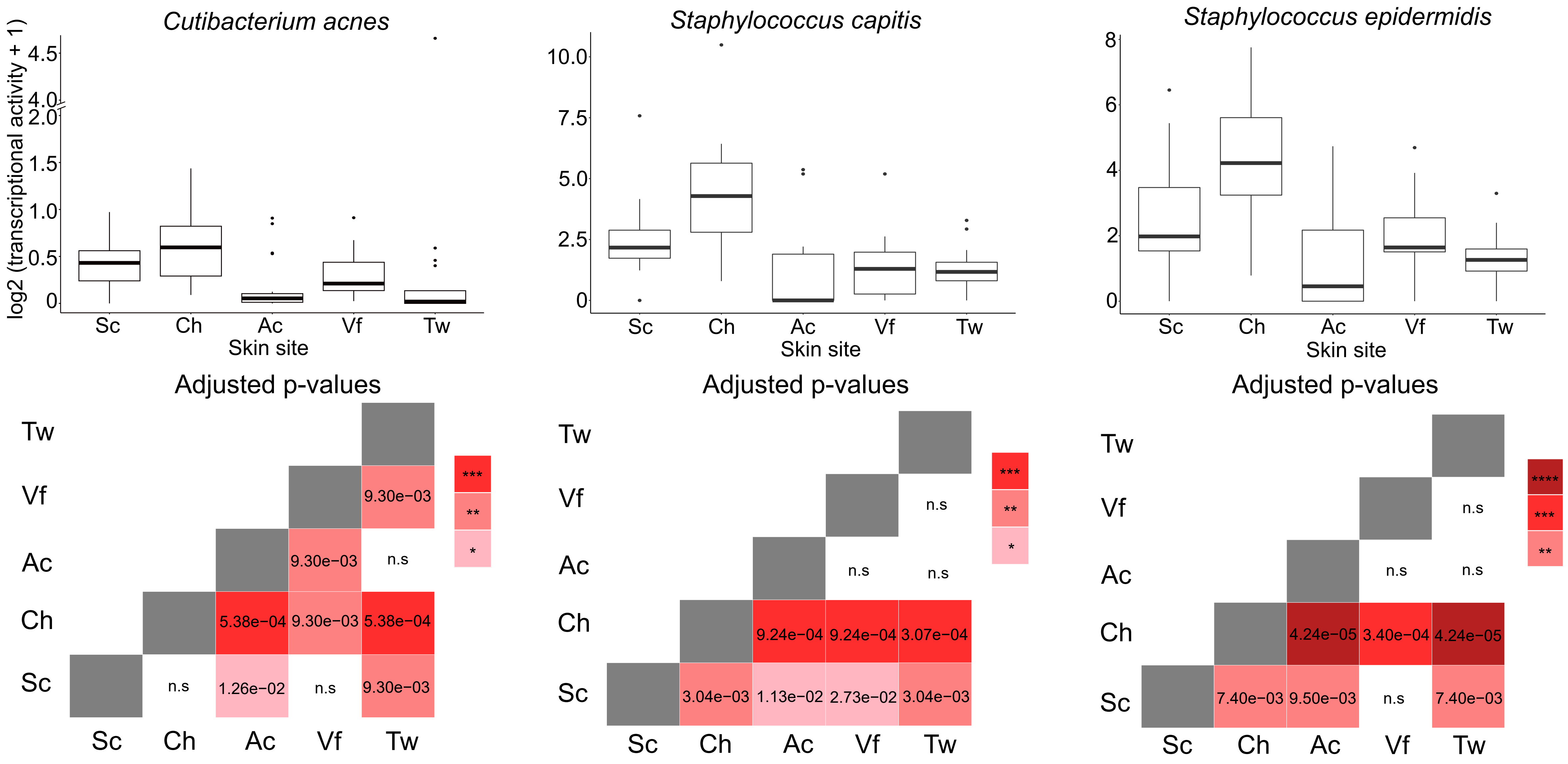


**Supplementary Figure 5: Transcriptional activity of selected prevalent bacteria across skin sites.** Boxplots of transcriptional activity (RNA/DNA) for each species, normalized by gene lengths and library sizes. Each site was represented by at least 5 libraries for which DNA reads for the given species were present at ≥0.1% relative abundance. Pairwise Wilcoxon rank sum tests were conducted and adjusted p-values are shown in the corresponding heatmaps. Data is from the full cohort (n=24, 23, 19, 18, 18 for scalp [Sc], cheek [Ch], antecubital fossa [Ac], volar forearm [Vf] and toe web [Tw] respectively).

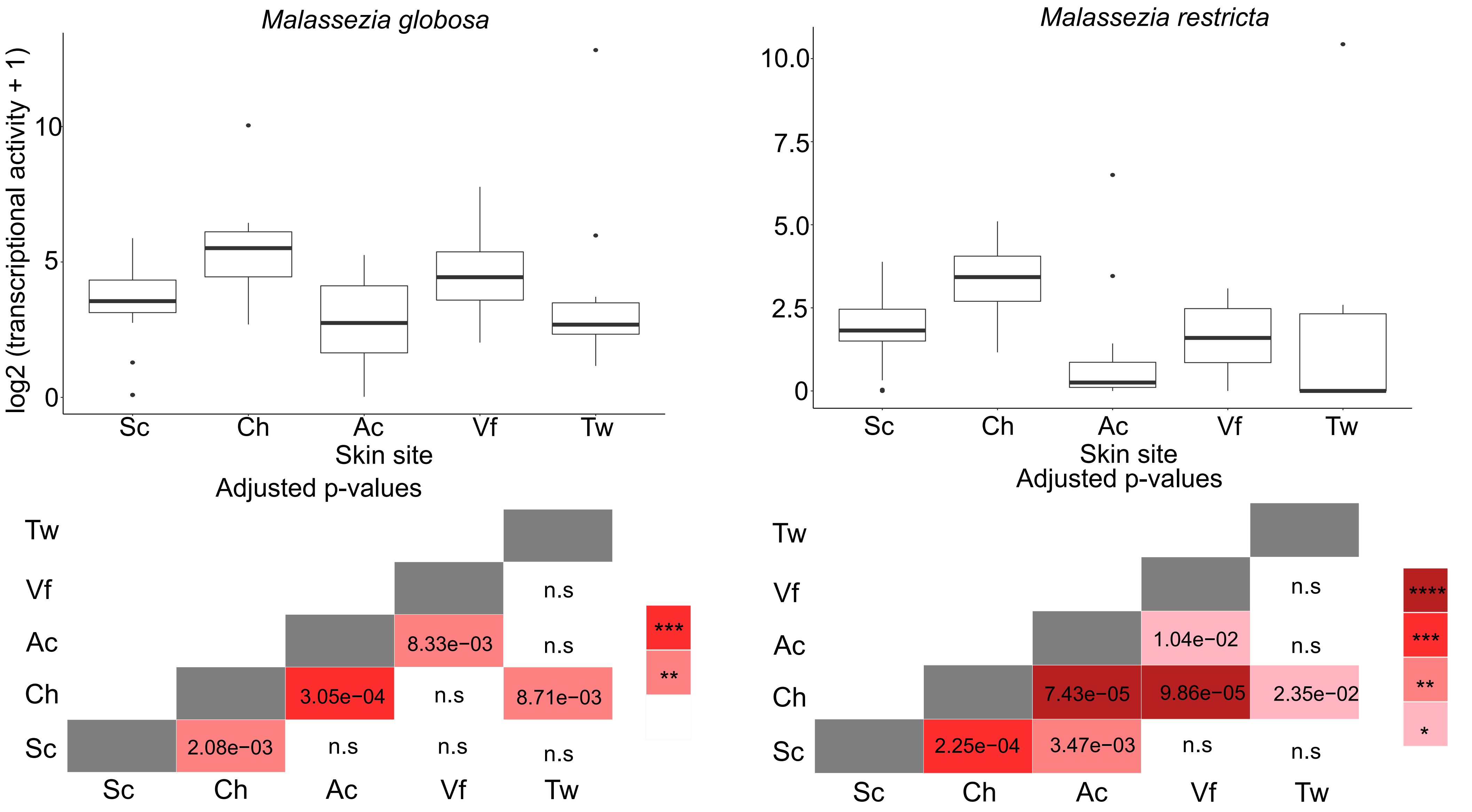


**Supplementary Figure 6: Transcriptional activity of selected prevalent fungi across skin sites.** Boxplots of transcriptional activity for each species (RNA/DNA), normalized by gene lengths and library sizes. Each site was represented by at least 5 libraries for which DNA reads for the given species were present at ≥0.1% relative abundance. Pairwise Wilcoxon rank sum tests were conducted and adjusted p-values are shown in the corresponding heatmaps. Data is from the full cohort (n=24, 23, 19, 18, 18 for scalp [Sc], cheek [Ch], antecubital fossa [Ac], volar forearm [Vf] and toe web [Tw] respectively).

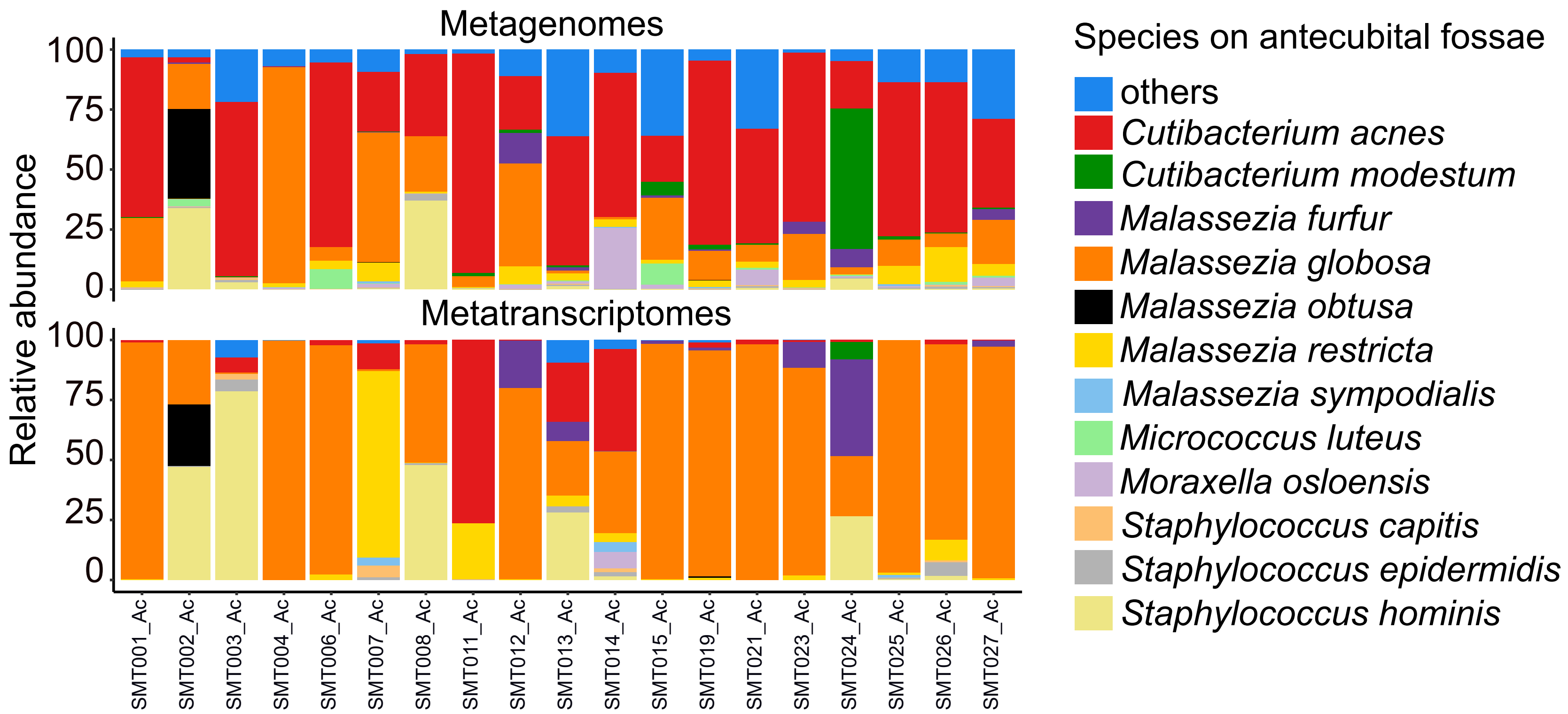


**Supplementary Figure 7: Comparison of metagenomic and metatranscriptomic species-level relative abundances in the antecubital fossae across individuals.** Stacked bar plots showing the most abundant species for matched skin metagenomes and metatranscriptomes across individuals (n=19).

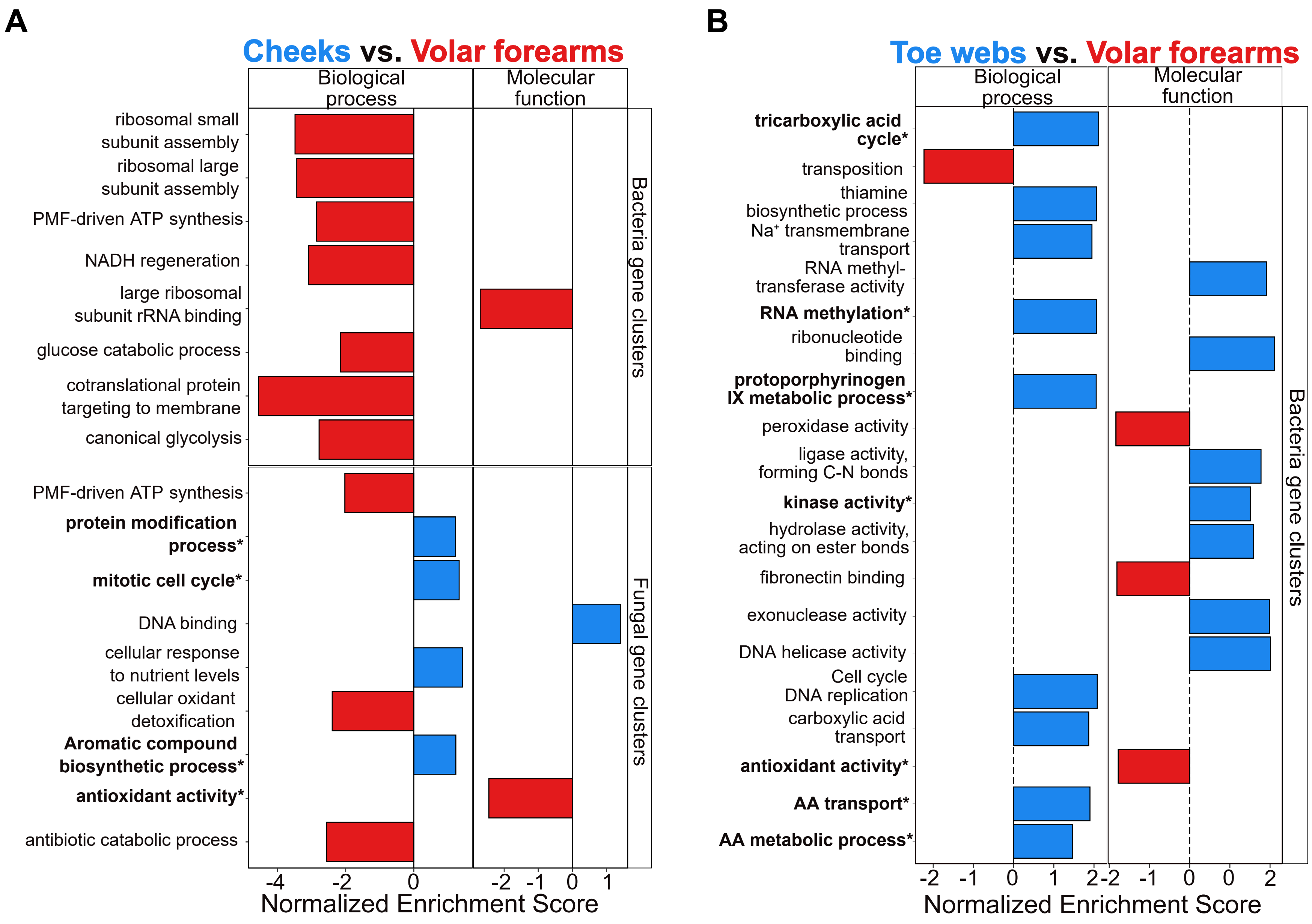


**Supplementary Figure 8: Functional enrichment of bacterial and fungal differentially expressed genes across skin sites.** (A) Barplots of normalized enrichment scores (NES) computed from gene set enrichment analyses of bacterial or fungal orthologous groups that are differentially enriched between cheek (n=22) and volar forearm (n=18) metatranscriptomes. (B) Same as (A) but computed between toe web (n=18) and volar forearm (n=18) metatranscriptomes. Biologically noteworthy pathways are bolded with an asterisk for clarity.

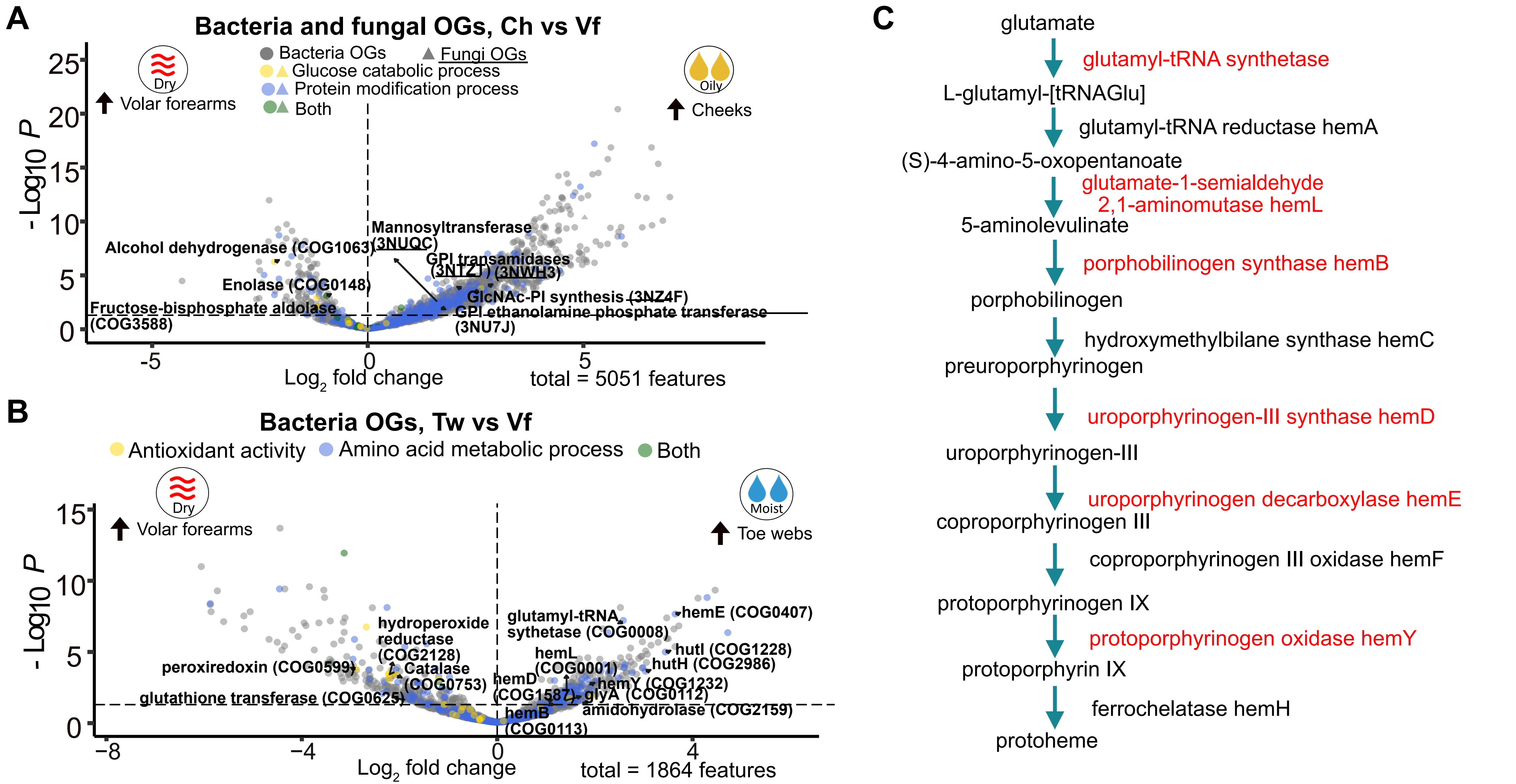


**Supplementary Figure 9: Differentially expressed microbial orthologous groups between different skin sites.** (A) Volcano plot of expressed bacterial and fungal orthologous groups (OGs) in cheeks (n=22) relative to volar forearms (n=18). (B) Volcano plot of expressed bacterial orthologous groups (OGs) in toe webs (n=18) relative to volar forearms (n=18). (C) Pathway map of heme biosynthesis from glutamate in prokaryotes. Enzymes highlighted in red were ≥2-fold upregulated in toe webs relative to volar forearm (adjusted p-value<0.05)

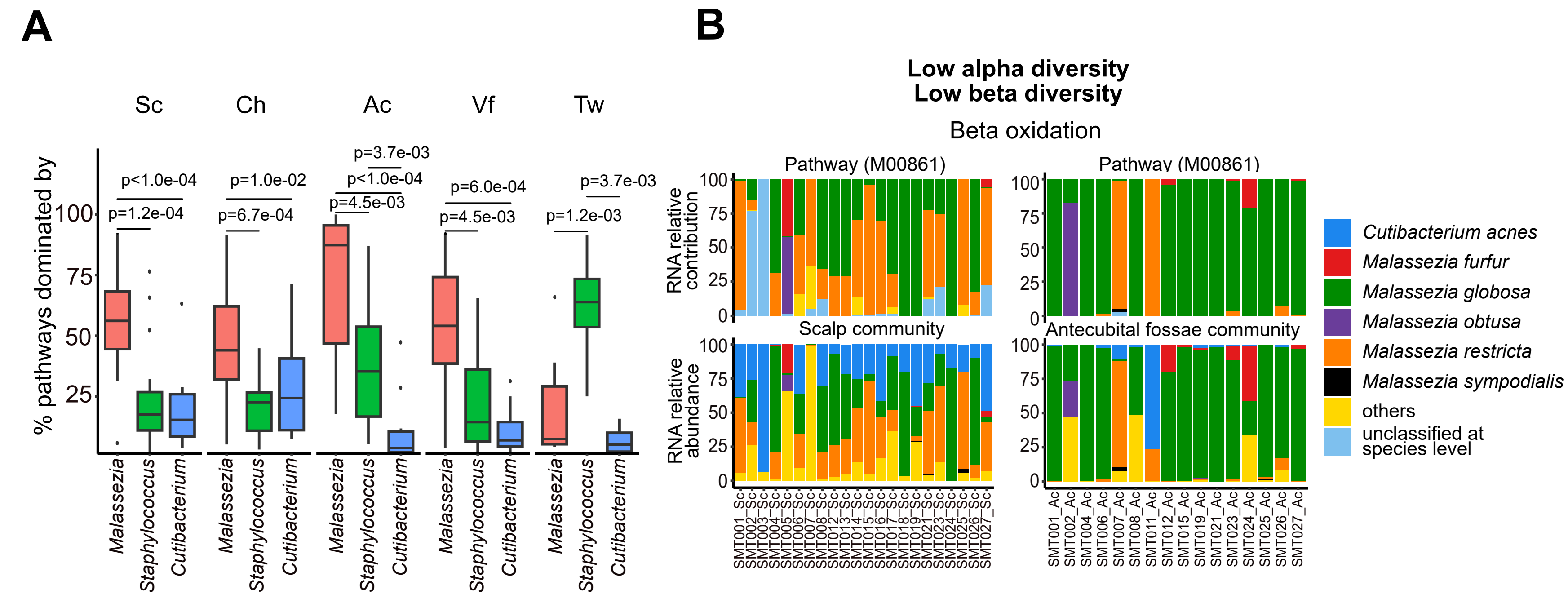


**Supplementary Figure 10: Pathway and community level contributions in skin metatranscriptomes.** (A) Boxplots of core microbial pathways dominated (>50% contribution) by *Malassezia*, *Staphylococcus* or *Cutibacterium*. Adjusted p-values for pairwise Wilcoxon ranked sum tests are shown. Stacked bar plots for species level pathway contributions at RNA level were estimated with HUMAnN3 for (B) beta oxidation of fatty acids, (C) galactose degradation by the Leloir pathway and (D) arginine biosynthesis. Community level relative abundances at RNA level were estimated with Kraken2.

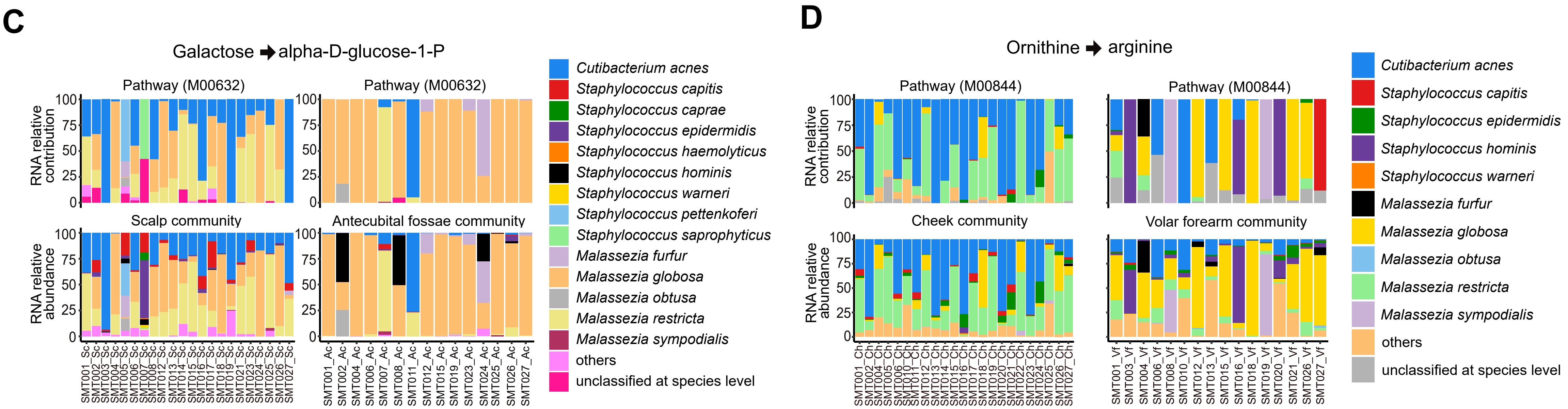

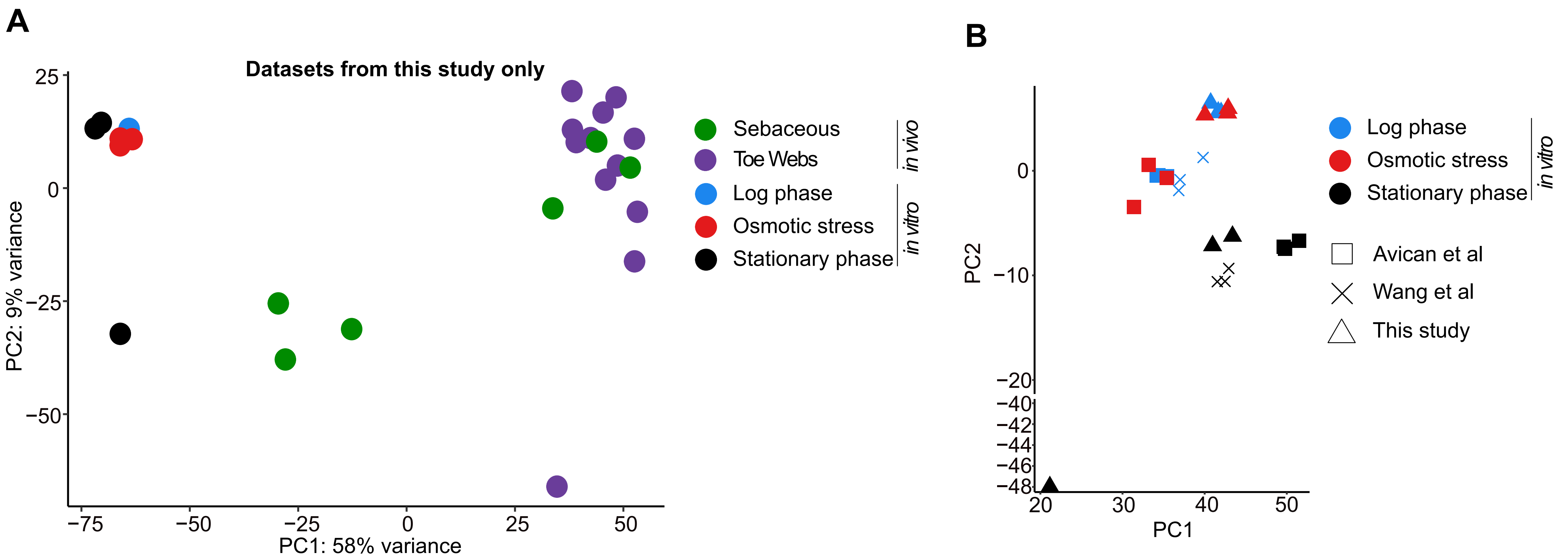


**Supplementary Figure 11: Comparison of *in vivo* and *in vitro* gene expression in *Staphylococcus epidermidis*.** (A) Principal component analysis (PCA) plot for *S. epidermidis* gene expression profiles from various *in vitro* and *in vivo* samples from the same experimental batch (this study only). (B) Principal component analysis (PCA) plot for *S. epidermidis* gene expression profiles from various *in vitro* conditions, keeping the same linear transformation as the PCA plot in **Figure 3E**. The different shapes represent three different experimental batches of *in vitro* samples. The y-axis is broken between -20 and -40 for visual clarity.

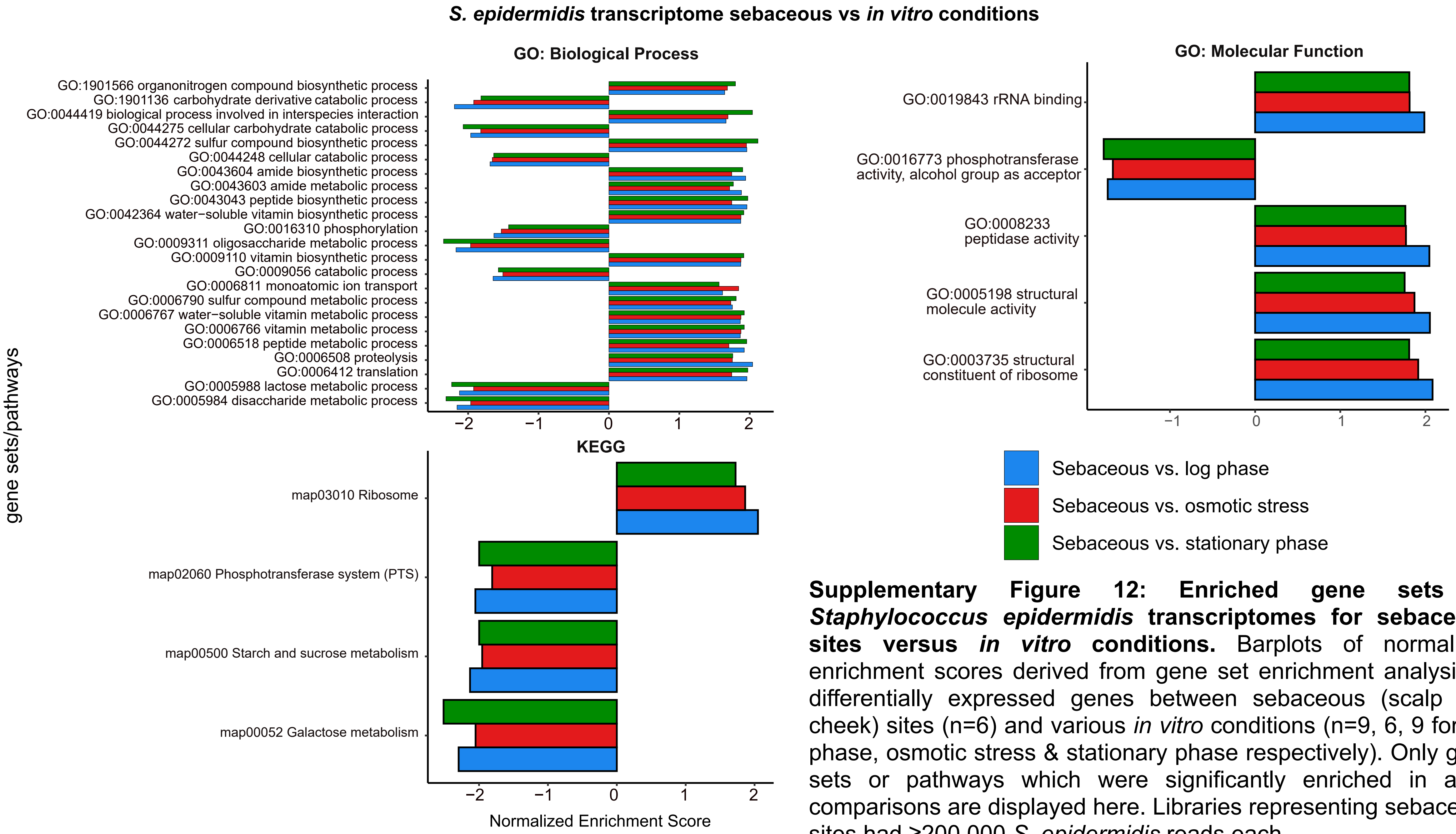


**Supplementary Figure 12: Enriched gene sets in *Staphylococcus epidermidis* transcriptomes for sebaceous sites versus *in vitro* conditions.** Barplots of normalized enrichment scores derived from gene set enrichment analysis of differentially expressed genes between sebaceous (scalp and cheek) sites (n=6) and various *in vitro* conditions (n=9, 6, 9 for log phase, osmotic stress & stationary phase respectively). Only gene sets or pathways which were significantly enriched in all 3 comparisons are displayed here. Libraries representing sebaceous sites had ≥200,000 *S. epidermidis* reads each.

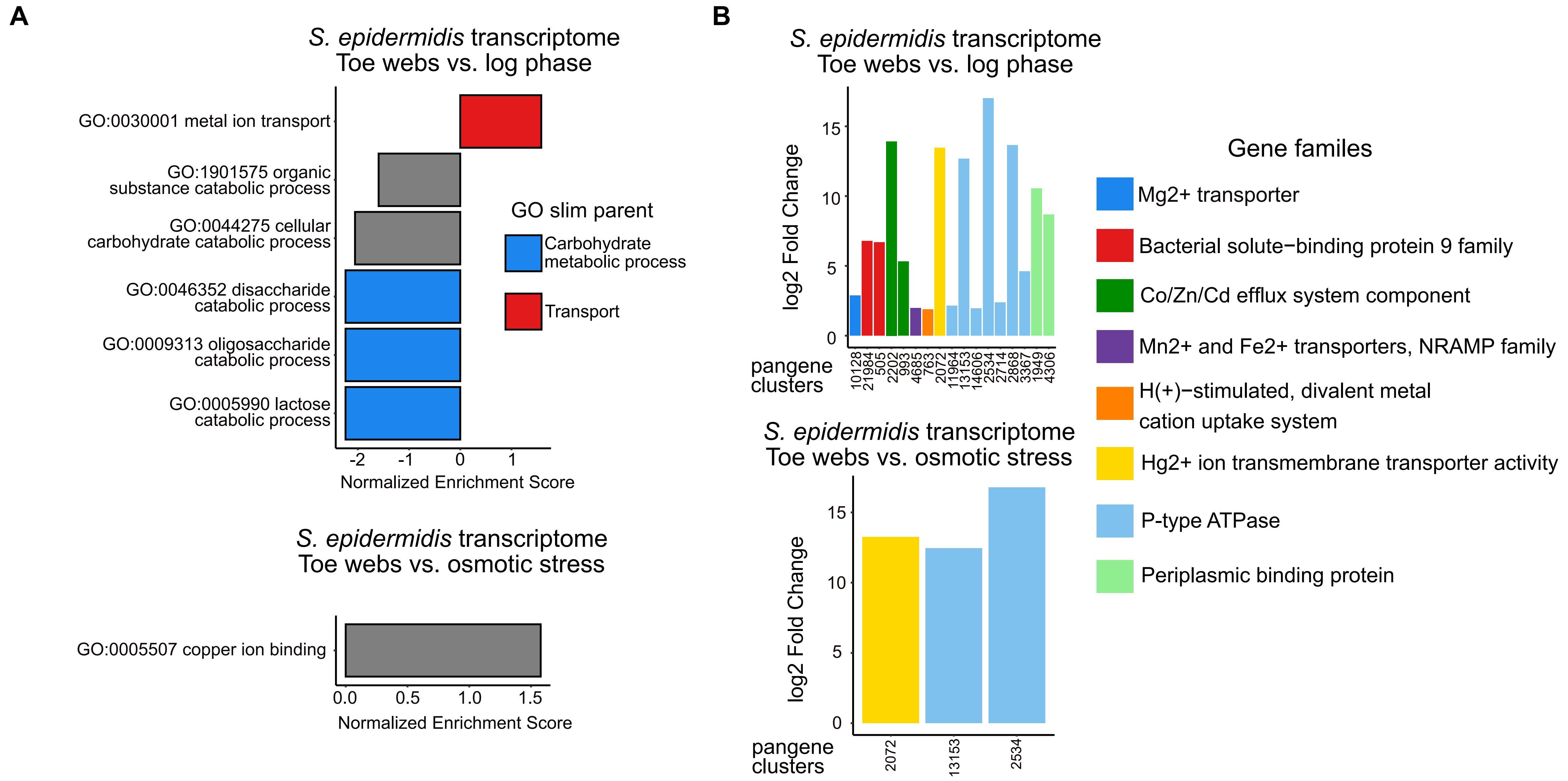


**Supplementary Figure 13: Enriched gene sets in *Staphylococcus epidermidis* transcriptomes for toe webs versus *in vitro* conditions.** (A) Barplots of normalized enrichment scores from gene set enrichment analysis between toe web sites (n=12) and various *in vitro* conditions (n=9, 6, 9 for log phase, osmotic stress & stationary phase respectively). (B) Barplots showing log2 fold change of genes related to metal ion transport (top) or copper ion binding (bottom), relative to *in vitro* conditions. Libraries representing toe webs had ≥200,000 *S. epidermidis* reads each.

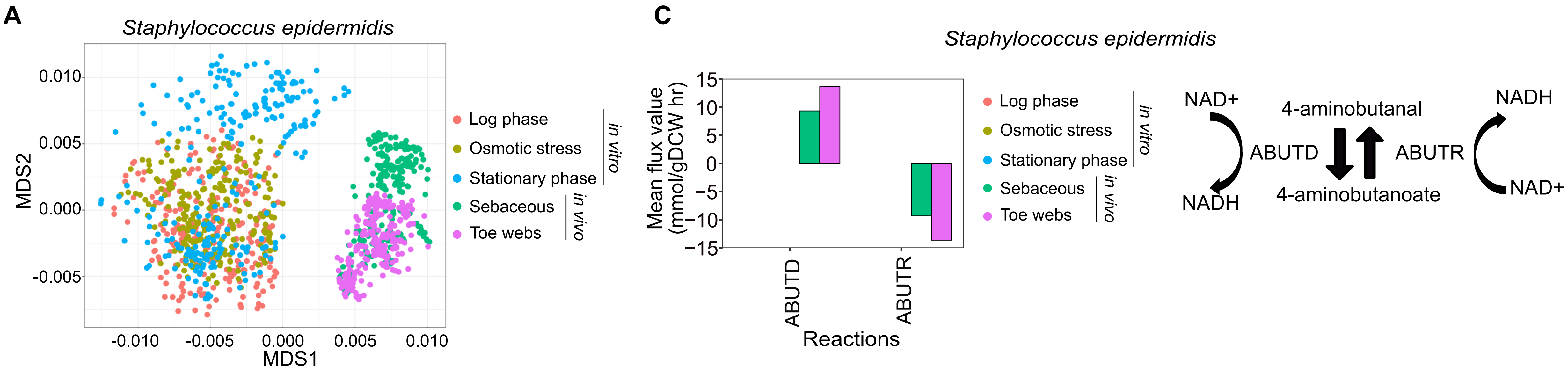


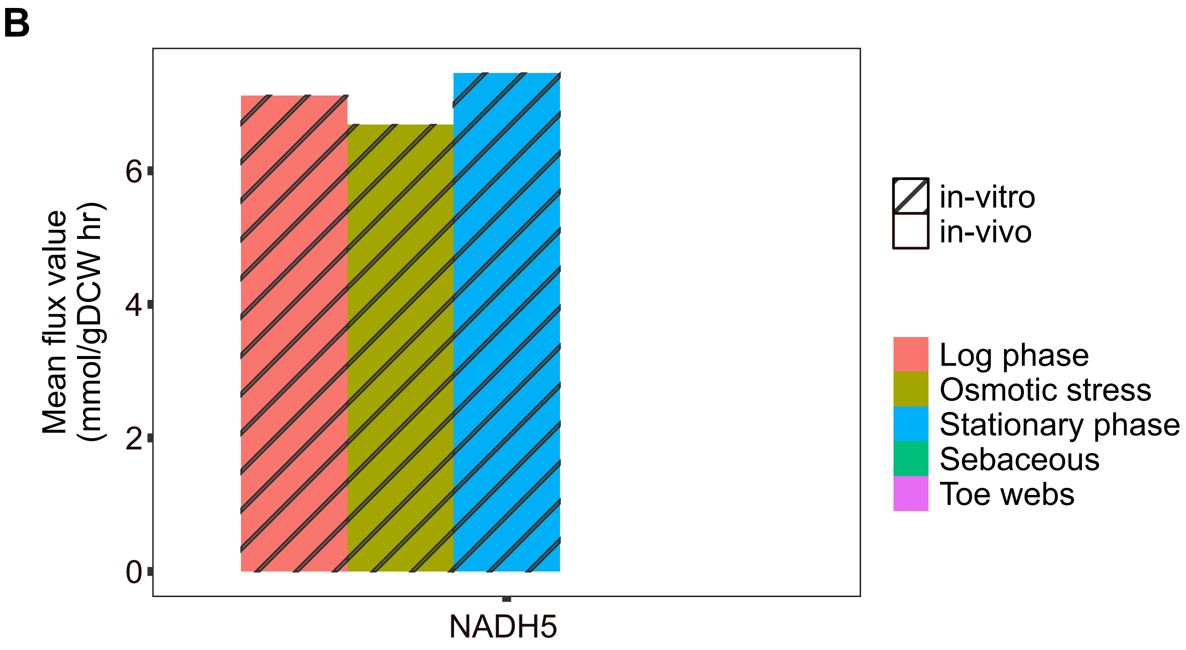


**Supplementary Figure 14: Comparison of metabolic fluxes for *Staphylococcus epidermidis* under *in vivo* and *in vitro* conditions***.* (A) MDS plot showing variation in metabolic flux between three *in vitro* (n=9, 6, 9 for log phase, osmotic stress & stationary phase respectively) and two *in vivo* (n=6, 12 for sebaceous sites and toe webs respectively) conditions. (B) Barplots showing mean flux values for the NADH5 reaction in *Staphylococcus epidermidis* which regenerates NAD+ via NADH dehydrogenase. (C) Barplots showing mean flux values for 4-aminobutyrate consumption (ABUTR) and production (ABUTD), which can also regenerate NAD+. Libraries for *in vivo* conditions had ≥200,000 *S. epidermidis* reads each.

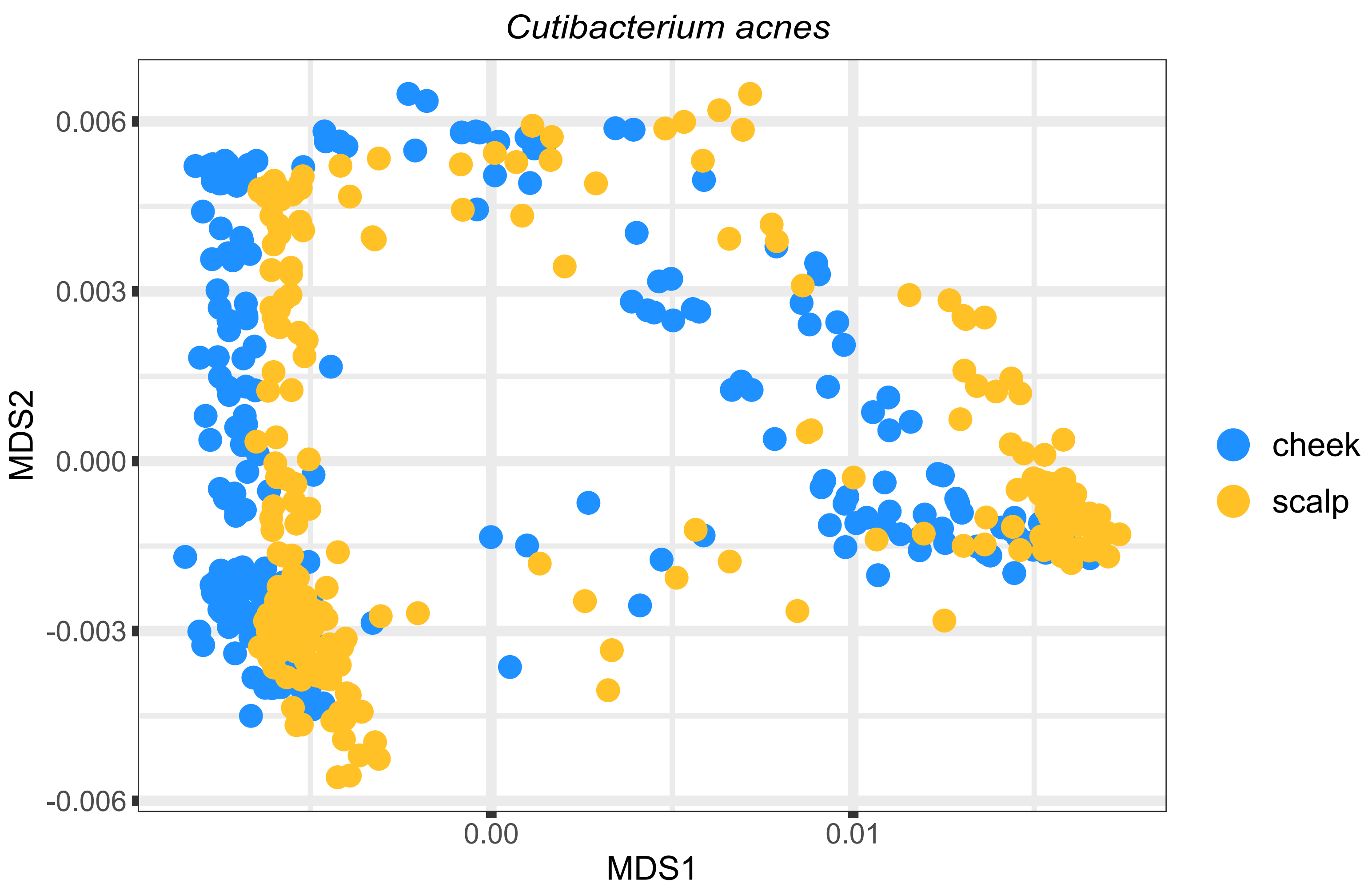


**Supplementary Figure 15: Comparison of genome-wide metabolic fluxes for *C. acnes* between cheek and scalp sites.** MDS plot showing variation in metabolic fluxes between two *in vivo* conditions: scalp (n=19) and cheeks (n=21) for *C. acnes.* Libraries for *in vivo* conditions had ≥200,000 *C. acnes* reads each.

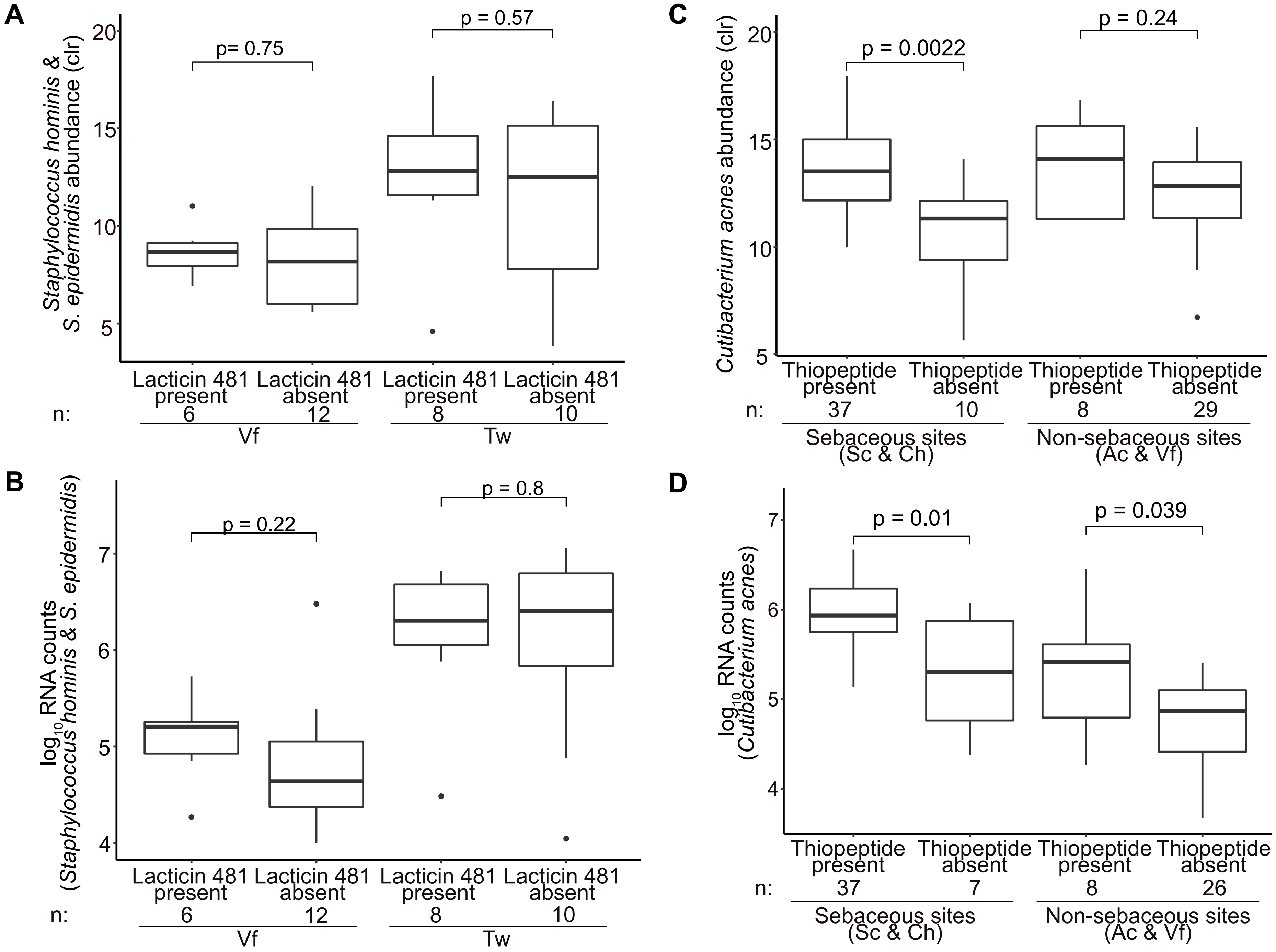


**Supplementary Figure 16: Variability of species and RNA abundances as a function of expression of selected antimicrobials.** (A) Boxplots of centered log ratio (CLR) DNA abundances of *S. hominis* & *S. epidermidis* from volar forearms or toe webs. Comparisons are between those with and without expression of lacticin 481 family peptides. (B) Same as (A), except showing the RNA counts for the species. (C) Boxplots of DNA abundances (CLR) of *C. acnes* from sebaceous or non-sebaceous sites, excluding toe webs. Comparisons are between those with and without expression of thiopeptides. (D) Same as (C), except showing the RNA counts for the species. Samples with no detected RNA counts for *C. acnes* were excluded. In all subfigures, the number of libraries in each category is given below its label (n).

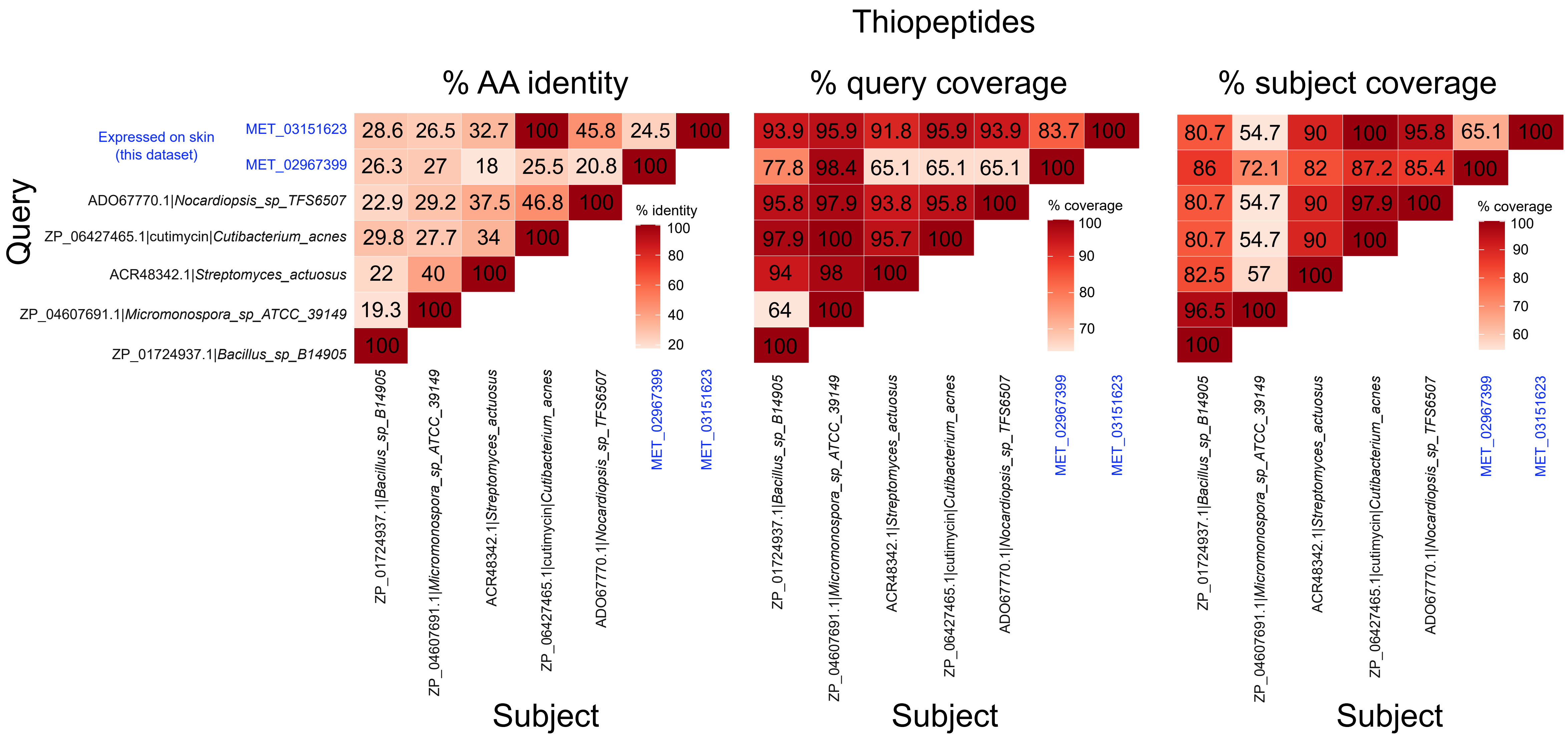


**Supplementary Figure 17: Percentage amino acid identity and coverage statistics from pairwise comparisons of thiopeptides.** Heatmaps showing various pairwise comparisons of thiopeptide amino acid sequences. Thiopeptide sequences were identified using profile hidden Markov models. Peptides highlighted in blue are expressed on skin in this dataset. Other peptides were chosen representatives from existing databases.

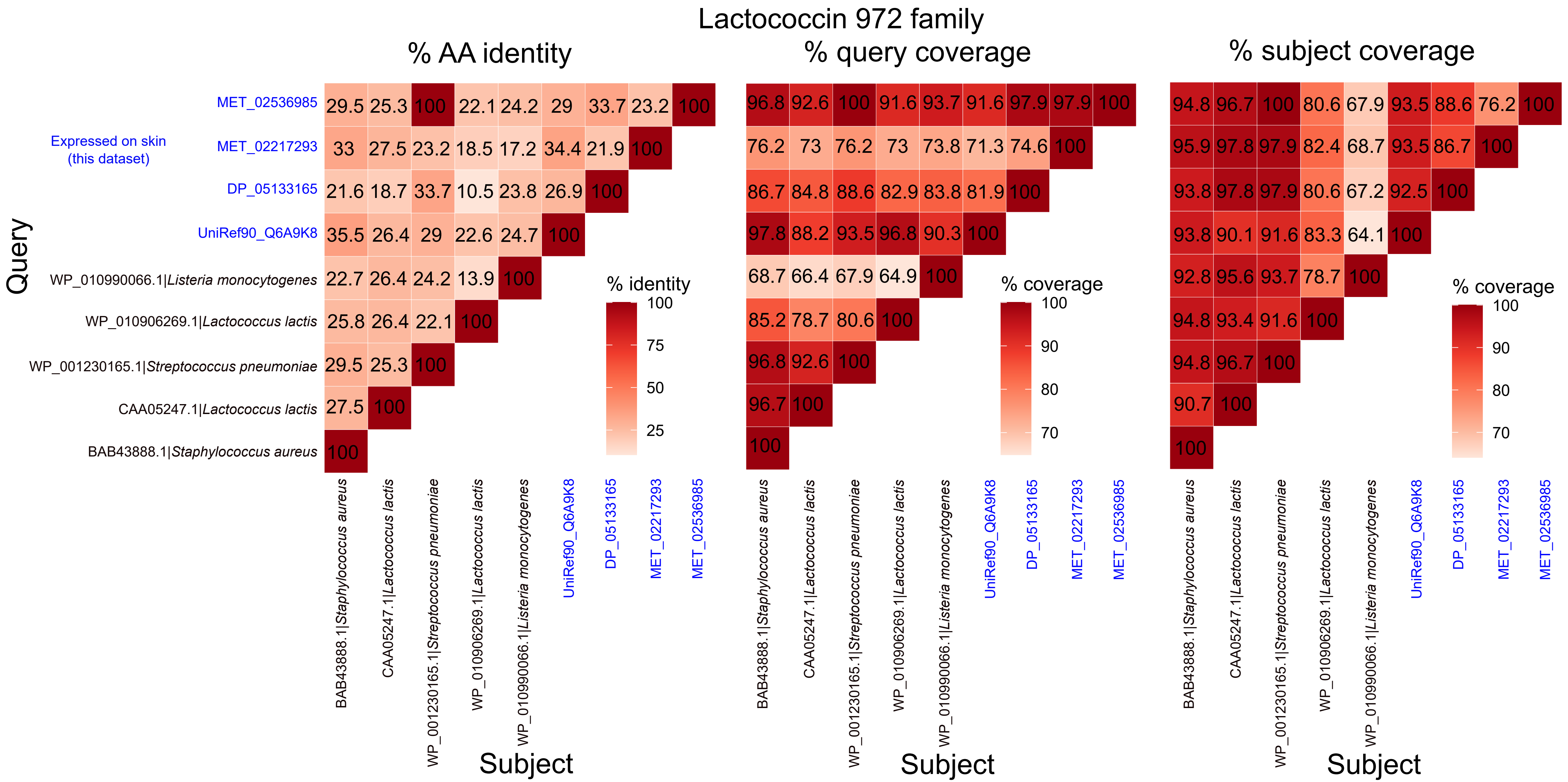


**Supplementary Figure 18: Percentage amino acid identity and coverage statistics from pairwise comparisons of bacteriocins of the lactococcin 972 family.** Heatmaps showing various pairwise comparisons of peptide sequences. Peptides of the lactococcin 972 family were identified using profile hidden Markov models. Peptides highlighted in blue are expressed on skin in this dataset. Other peptides were chosen representatives from existing databases.

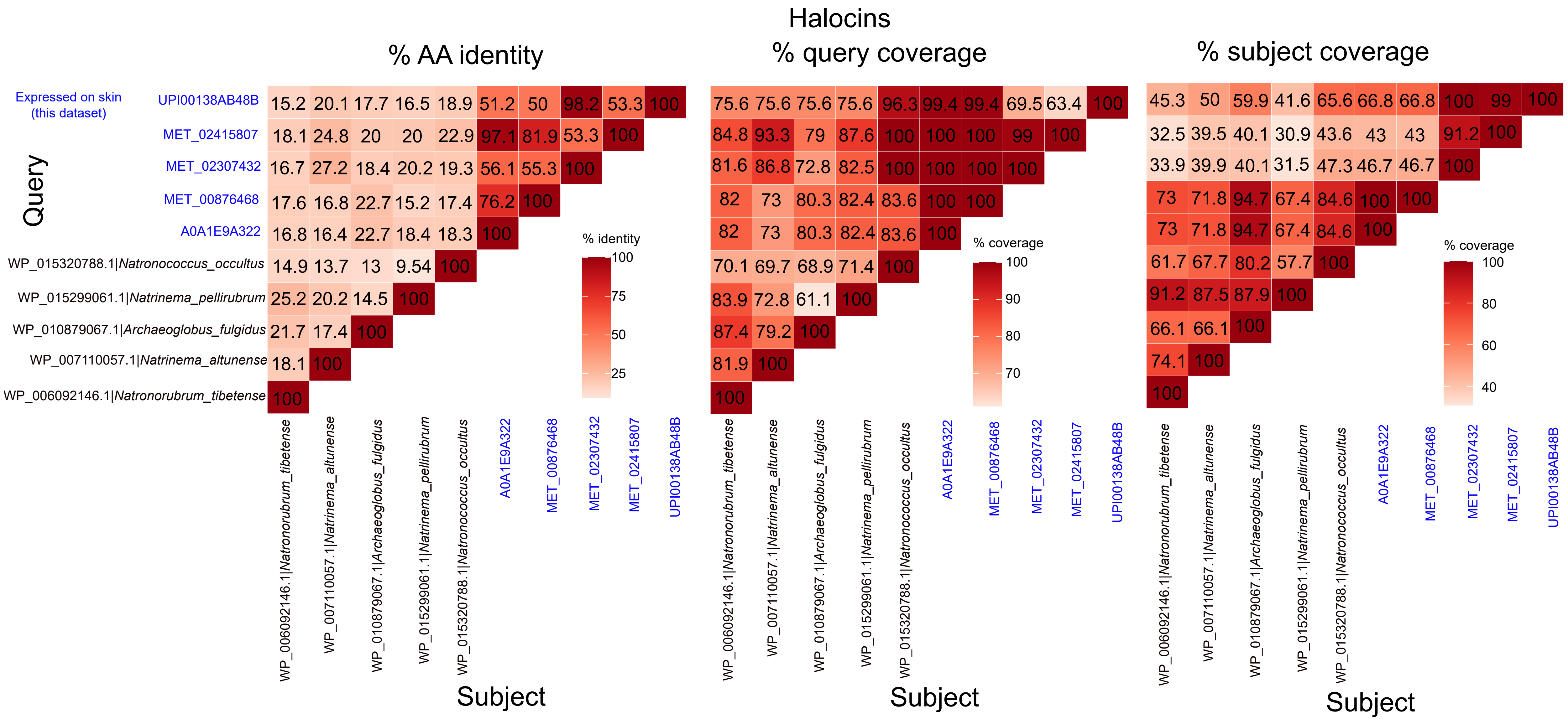


**Supplementary Figure 19: Percentage amino acid identity and coverage statistics from pairwise comparisons of halocins.** Heatmaps showing various pairwise comparisons of peptide sequences. Peptides of the halocin family were identified using profile hidden Markov models. Peptides highlighted in blue are expressed on skin in this dataset. Other peptides were chosen representatives from existing databases.

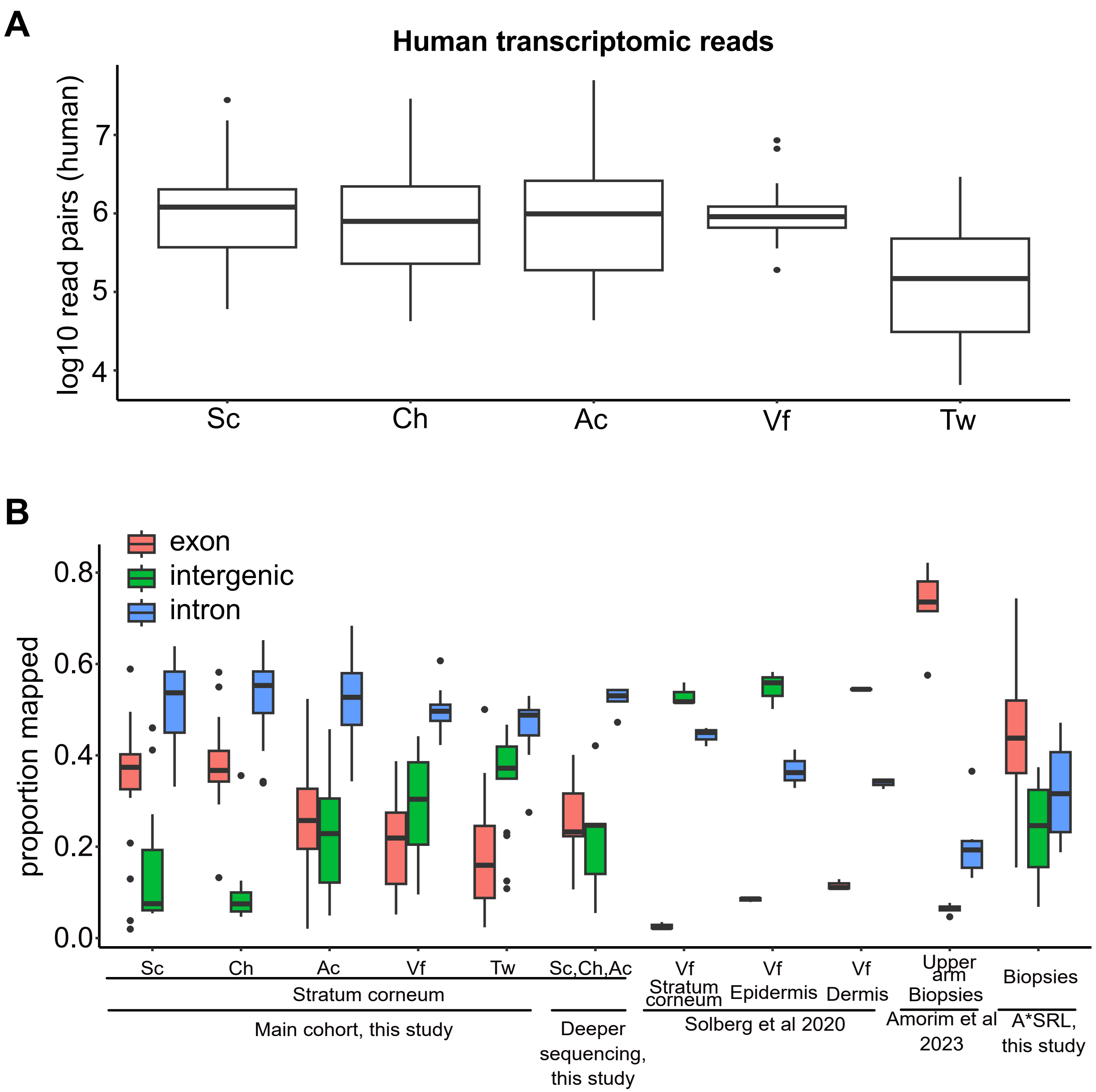


**Supplementary Figure 20: Quality control metrics for human reads in skin metatranscriptomic libraries.** (A) Boxplots depicting number of sequenced human reads across different skin sites from this study. (B) Boxplots showing the proportion of human RNA reads mapping to exons, intergenic regions and introns. The full cohort for this study comprised of n=24, 23, 19, 18, 18 libraries for scalp (Sc), cheek (Ch), antecubital fossa (Ac), volar forearm (Vf) and toe web (Tw) respectively. The samples in this study subjected to deeper re-sequencing comprised of n=2, 3, 3 libraries for Sc, Ch and Ac respectively. The samples for Solberg et al 2020 comprised n=3 libraries each for the stratum corneum, epidermidis and dermis. The samples for Amorim et al comprised n=6 libraries for upper arm biopsies. The biopsies from A*SRL comprised n=4 libraries.

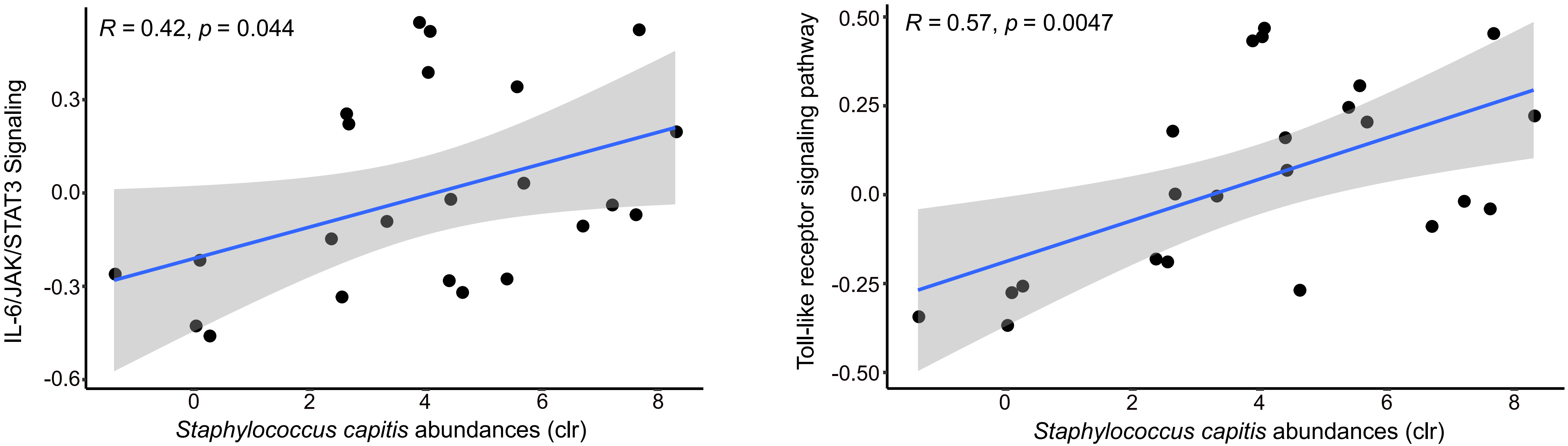


**Supplementary Figure 21: Correlations between estimated human pathway activity and *Staphylococcus capitis* abundances.** Scatterplots showing positive correlations between immune pathway activity estimated by gene set variation analysis (GSVA) and centered log ratio transformed abundances of *Staphylococcus capitis* on cheeks.

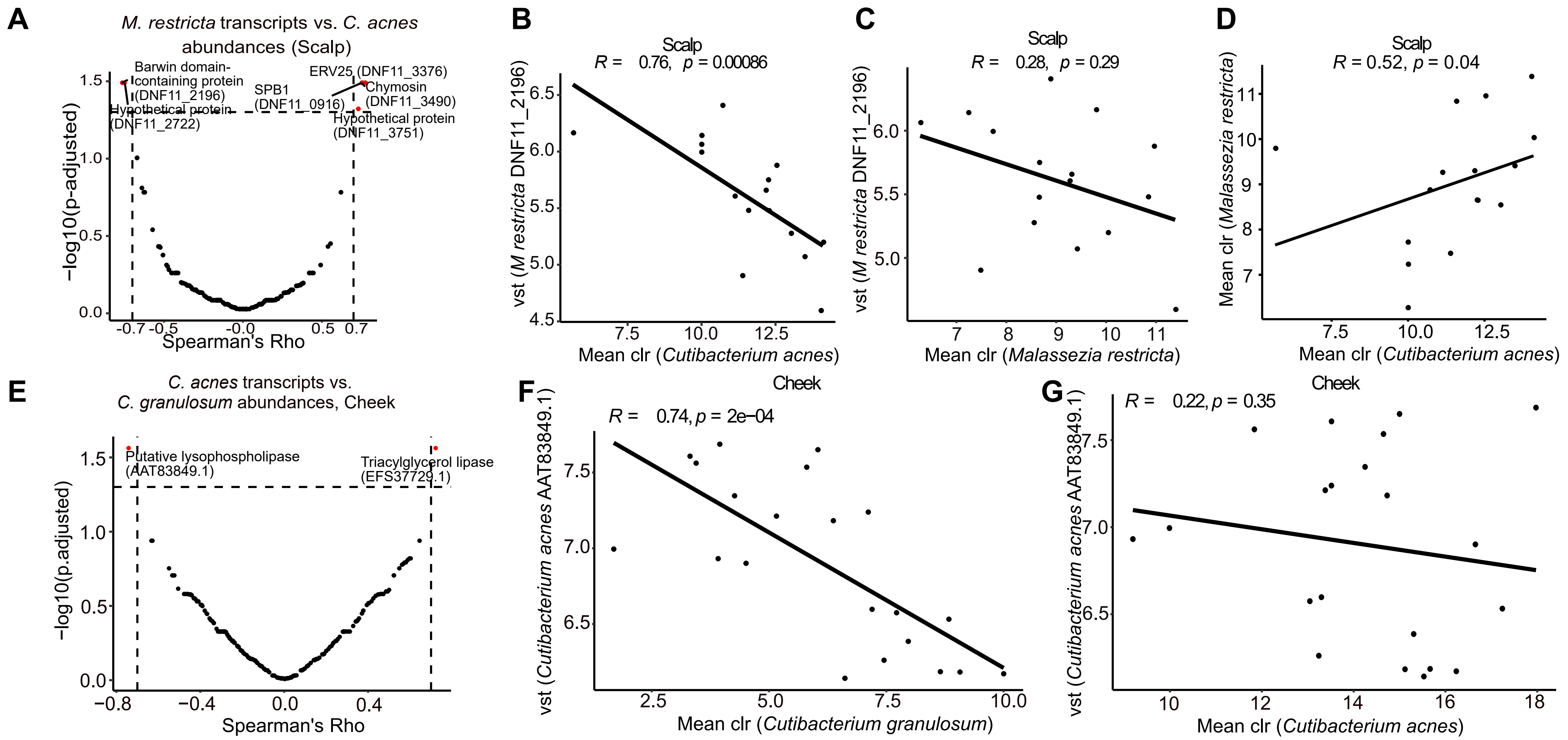


**Supplementary Figure 22: Correlations between microbial transcript and organism abundances of two distinct species on the same site.** (A) Scatterplot of adjusted p-values and Spearman Rho coefficients derived from correlating *Malassezia restricta* transcript abundances with *Cutibacterium acnes* metagenomic abundances across individuals on scalp sites.(B) Scatterplot showing negative correlation between variance stabilized (vst) levels of a transcript from a Barwin domain-containing protein (DNF11_2196) expressed in *Malassezia restricta* and centered log ratio (clr) abundances of *Cutibacterium acnes.* (C) Same as (B) but showing the relationship between the transcript and clr abundances of *M. restricta*. (D) Scatterplot of clr abundances of *M. restricta* against *C. acnes*. (E) Volcano plot showing correlation and adjusted *p*-values of transcripts of *C. acnes* proteins in the secretory pathway against clr abundances of *Cutibacterium granulosum.* Statistically significant correlations are highlighted in red. (F) Scatterplot showing negative correlation between levels of a transcript (AAT83849.1) expressed in *C. acnes* and clr abundances of *C. granulosum*. (G) Same as (F) but showing the relationship between the transcript and clr abundances of *C. acnes.*

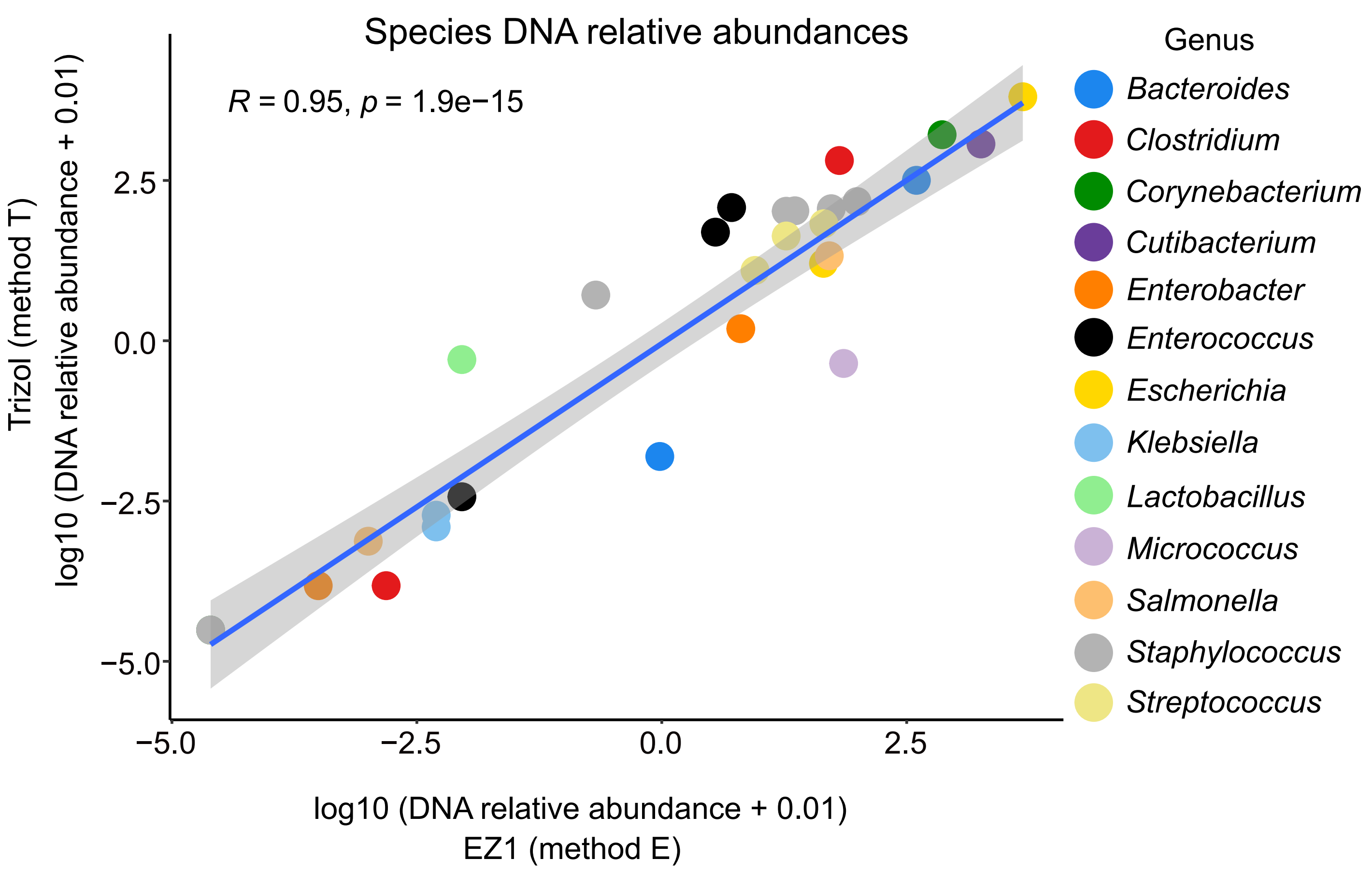


**Supplementary figure 23: Correlations between microbial species-level DNA abundances from mock communities extracted using two different methods.** Scatterplot comparing relative abundance profiles for microbial DNA extracted with a trizol method (method T) versus DNA extracted using a standard Qiagen kit (method E). Results are from three different mock communities of common human microbes. Taxonomic profiles were obtained using Kraken 2 with default parameters. The Pearson correlation coefficient and corresponding p-value are shown in the figure.
